# Current protein structure predictors do not produce meaningful folding pathways

**DOI:** 10.1101/2021.09.20.461137

**Authors:** Carlos Outeiral, Daniel A. Nissley, Charlotte M. Deane

## Abstract

Protein structure prediction has long been considered a gateway problem for understanding protein folding. Recent advances in deep learning have achieved unprecedented success at predicting a protein’s crystal structure, but whether this achievement relates to a better modelling of the folding process remains an open question. In this work, we compare the pathways generated by state-of-the-art protein structure prediction methods to experimental folding data. The methods considered were AlphaFold 2, RoseTTAFold, trRosetta, RaptorX, DMPfold, EVfold, SAINT2 and Rosetta. We find evidence that their simulated dynamics capture some information about the folding pathwhay, but their predictive ability is worse than a trivial classifier using sequence-agnostic features like chain length. The folding trajectories produced are also uncorrelated with parameters such as intermediate structures and the folding rate constant. These results suggest that recent advances in protein structure prediction do not yet provide an enhanced understanding of the principles underpinning protein folding.

## 1. INTRODUCTION

Protein folding, or how a protein attains its equilibrium three-dimensional structure, is considered one of the grand challenges of modern molecular biology (1). If it were possible to accurately predict the folding pathway of a protein, it would have far-reaching implications for basic science, further the development of novel therapeutics and broaden the toolset for protein design and engineering. Some of the most prevalent aging-related pathologies, like Alzheimer’s (2) or Parkinson’s disease (3), originate when the delicate proteostasis machinery fails to ensure that proteins are correctly folded. The dynamical nature of the folding process also relates to other poorly understood phenomena like allostery (4), fold-switching (5) or intrinsic disorder (6). Even protein expression, one of the cornerstones of modern biotechnology, is highly dependent on folding: problems expressing recombinant proteins across different organisms are often attributed to changes in the folding mechanism due to different translation machinery (7). However, we are still unable to accurately predict the folding pathway of a protein *de novo*.

The related problem of protein structure prediction has experienced significant progress over the past two decades, powered by the community-wide effort of the biennial CASP contest (8). This assessment exercise has witnessed multiple step changes in accuracy as novel ideas have been incorporated into the participant’s pipelines (9; 10; 11). In recent years, deep learning approaches have dramatically improved the quality of structure prediction. The introduction of deep learning techniques into protein structure prediction methods raised the average free modelling GDT_TS score, which measures structural similarity on a scale from 0 to 100, from 52.9 in CASP12 (10), to 65.7 in CASP13 (11). In CASP14, a deep learning model, AlphaFold 2, achieved an average GDT_TS of 85.1 (12). This method, and other similar techniques (13), have been hailed as an acceptable solution to the protein structure prediction problem (14).

These dramatic advances in structure prediction raise a fundamental question: has progress been driven by a better understanding of protein folding? To the best of our knowledge, the ability of structure predictors to determine folding pathways has not been evaluated previously. Related work has studied the search trajectories of fragment replacement methods (15), or attempted to introduce biological constraints into folding (16). Furthermore, recent work has shown that some deep learning predictors can pinpoint flexible residues (17) or conformational changes (18), suggesting that these methods may capture dynamic phenomena encoded in the multiple sequence alignment. In this work we examine whether protein structure prediction methods are able to reveal anything about a protein’s folding pathway.

**Figure 1.**
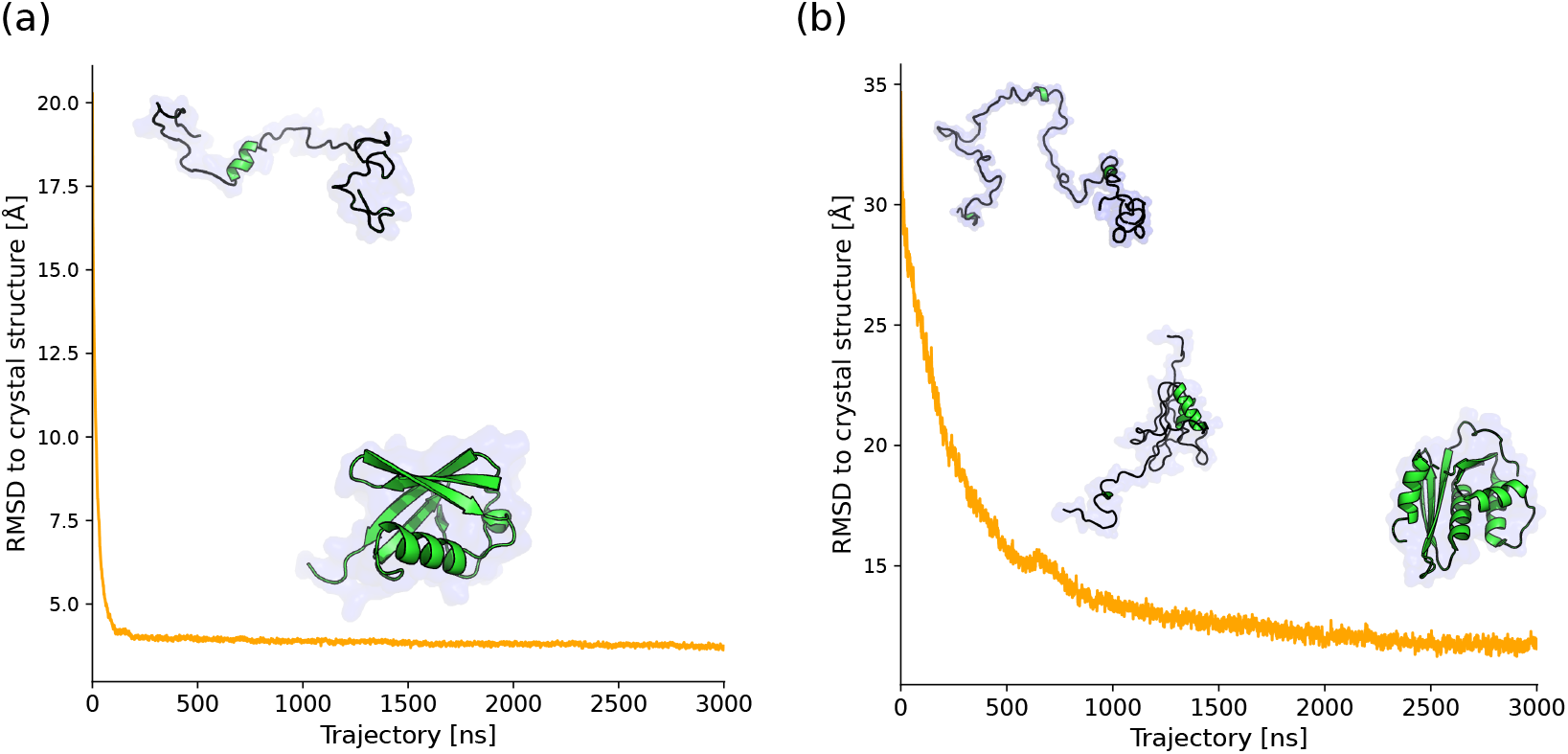
Two-state vs multistate protein folding pathways. Some proteins fold in a single cooperative step (two-state folding, in reference to the two states: unfolded and folded states), while others exhibit complex pathways with one or more intermediates (multistate). In the plots, we show the average of 200 refolding coarse-grained molecular dynamics simulations, superposed with representative snapshots of the structure. (a) Human ubiquitin (PDB: 1UBQ), a two-state folder: the protein folds immediately to a structure that resembles the crystal structure. (b) The RNAse H domain of HIV-1 reverse transcriptase (PDB: 1HRH), which folds through a stable intermediate where the majority of the protein is compact, except for the C-terminal α-helix that is unstructured.

We show that current protein structure prediction methods do not produce correct folding pathways. We first demonstrate that generated pathways have a weak link to formal folding kinetics, achieving a modest accuracy in discerning between protein chains that fold in a two-state or multistate mechanism. However, a simple sequence-agnostic feature, the length of the protein chain, is a far better predictor of folding dynamics. In the case of two-state folders, we also find that the dynamic trajectory is inconsistent with experimental folding rate constants. We then demonstrate that predicted pathways produce erratic intermediates that are inconsistent with available hydrogen-deuterium exchange data. Most of the structure prediction methods are not significantly better than an unbiased coin and some of them are consistently worse at reproducing experimental measurements. Finally, we take a protein with well-characterised folding pathway, ubiquitin, and compare experimental data, simulations and pathways derived from protein structure prediction,

## 2. METHODS

### 2.1. Reference data

We compiled a dataset of 170 proteins for which experimental folding kinetics data is available. To produce this dataset, we collated entries from the Protein Folding Database (PFDB) of kinetic constants (19) and the Start2Fold directory of hydrogen-deuterium exchange experiments (20). We checked the annotations contained in the PFDB and changed the classification for human ubiquitin (PDB: 1UBQ) from multistate to two-state, given that the PFDB citation corresponds to a mutated species and the wild-type protein displays two-state kinetics (21). The entries in the Start2Fold database do not include annotation for formal kinetics, so we manually annotated the results by querying the literature. The complete dataset and original publications are provided in Appendix A. We also compiled folding rate constants for a fraction of the proteins in this dataset that exhibit two-state kinetics, which are reported in Appendix B.

We collected available hydrogen-deuterium exchange (HDX) data from Start2Fold and original papers (Appendix C), to use as structural insight into the folding pathway. We observed that the residue-level annotation in the original database was sparse; we therefore queried the original sources and reconstructed the annotation as indicated in Appendix C. Each secondary structure element was labeled as structured or unstructured for each of the identified intermediates, on the basis of the experimental protection factors of the probes (in NMR experiments) or peptides (in mass spectrometry experiments) corresponding to a given portion of secondary structure.

**Figure 2.**
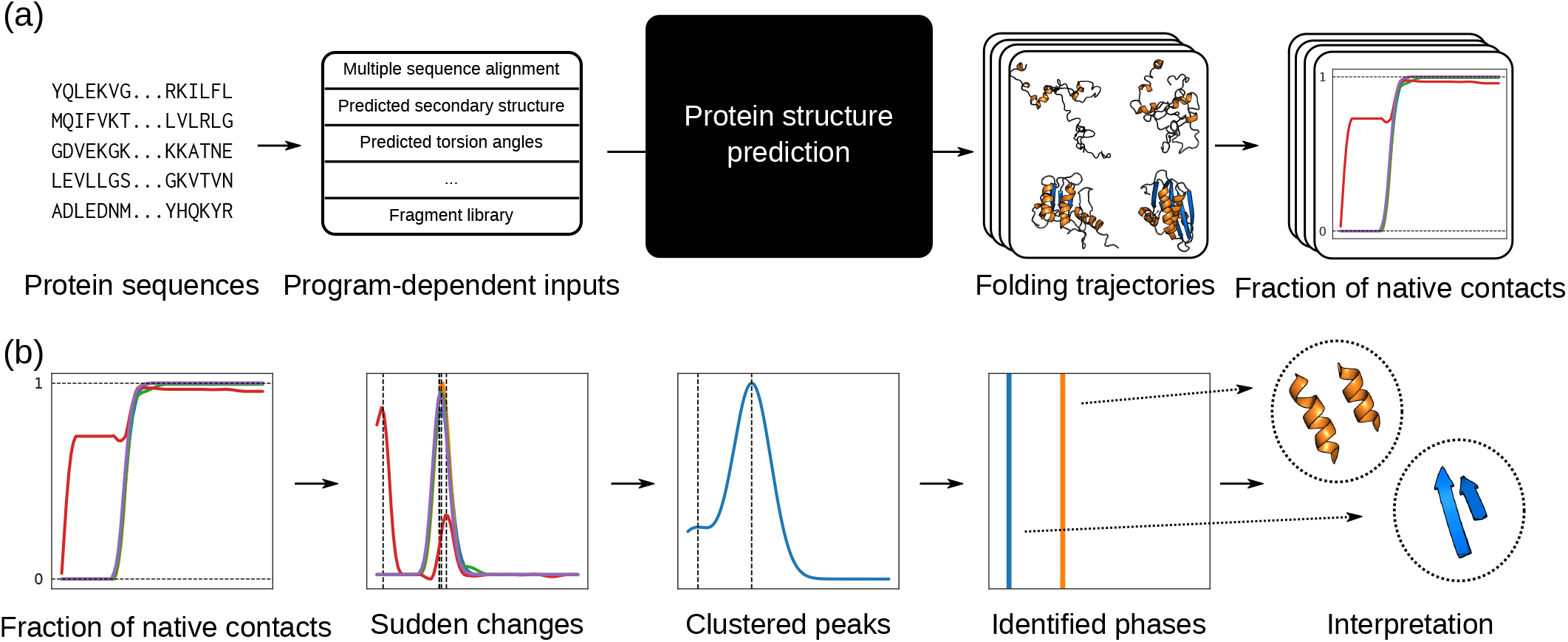
Protocol for the analysis of simulated folding pathways. (a) Trajectory generation process. Protein sequences are used to generate the necessary input features for a modified protein structure predictor using default processing scripts. The structure prediction software outputs detailed search trajectories, that are then summarised as the fraction of native contacts between pairs of secondary structure elements. (b) The trajectories are smoothed, and the positions of maximum change are identified via numerical differentiation. These peaks are subsequently clustered using kernel density estimation (KDE) with a Gaussian kernel, allowing us to identify main phases of folding, and establishing whether the trajectory proceeds in one or more steps; and into the structural intermediates, which can be compared to hydrogen-deuterium exchange (HDX) experiments.

Sequences and reference structures were downloaded from the RCSB PDB (22) and trimmed according to the specifications of the entries. We used the codes referenced in the publications, even when higher resolution structures were available in the PDB. When using NMR structures with multiple models, the structure with the highest score was selected. Missing regions were repaired using MODELER (23) with standard parameters.

### 2.2. Trajectory generation

We generated protein folding trajectories using the latest versions, as of December 2020, of Rosetta (24), trRosetta (25), DMPfold (26), EVcouplings (27), RaptorX (28), SAINT2 (16), and the recently published RoseTTAFold (13). We modified the source codes of the seven programs to print the current structure after every fragment substitution (for Rosetta and SAINT2); or after every 10 gradient updates (for trRosetta, RaptorX, DMPfold and EVfold, which employ L-BFGS or related gradient descent algorithms); or after every refinement cycle in a SE(3)-equivariant iterative transformer (for RoseTTAFold). Given the large amount of data produced by Rosetta, averaging more than 200,000 snapshots per decoy, we subsampled the trajectories produced at every 100 fragment substitutions.

We preprocessed the sequences of our 170 test case proteins employing the default pipelines provided by each piece of software, and used default parameters throughout. The generated trajectories for each of the 170 annotated proteins were compressed to the binary DCD format (29) and analysed using in-house scripts. For RoseTTAFold, which produces only the atoms involved in the peptide bond, we used PULCHRA (30) to reconstruct the *β*-carbons which are used in subsequent analysis. All information necessary to reproduce this study, including the `diff` files of the original source code, is available from.

We also considered trajectories generated by AlphaFold 2 (14). Due to the architecture of the model, producing a trajectory would require training a replica of the AlphaFold Structure Module for every individual Evoformer iteration; this was done by Jumper *et al*. in the original publication, although the models have not been open-sourced. Fortunately, individual folding trajectories for each of the 170 proteins in our dataset were kindly provided by the DeepMind team. These trajectories were generated with the same methods and models as in the original publication (14), save for the removal of any templates (although, of course, many of the structures were present in the training set).

### 2.3. Trajectory analysis

We analysed the trajectories using the fraction of native contacts between secondary structure elements (31). These elements were identified using STRIDE (32) on the crystal structure, ignoring any element shorter than 4 amino acids. Distances were calculated using MDAnalysis (33; 34), and two amino acids were defined to be in contact if their *β*-carbons (*α*-carbons in the case of glycine) were less than 8.0 *Å* apart in the native structure. To account for fluctuations, we introduced a flexibility parameter *ξ* = 1.2 whereby amino acids in contact in the crystal structure were still considered to be in contact in the simulated trajectory if their distance was *ξ* times the crystal structure distance. These parameter choices were inspired by the standard in the molecular dynamics literature *e.g*. (35). To ensure that our conclusions were independent of the choice of parameters, we performed a parameter exploration on a reduced subset of the data (10 trajectories per protein) – see Figure S1.

We computed the numerical time derivatives of the fraction of native contacts using finite differences and smoothed them using Friedman’s supersmoother (36) as implemented in the R `stats` package (37). The maximum value of the derivative for a pair of secondary structure elements was identified as the time point where the two of them are folded. We then fitted the data using a Gaussian Kernel Density Estimation (KDE) with bandwidth determined by Scott’s rule via SciPy (38). When all of the folding transitions belong to a single peak, the trajectory was considered to be folding in two-states; when two or more peaks were found, the trajectory was labeled as multistate. Given the variability of the trajectories between prediction runs, many proteins had both two-state and multistate trajectories; hence we defined the fraction of two-state trajectories as the probability that a protein exhibits two-state kinetics.

### 2.4. Coarse-grained molecular dynamics simulations

Human ubiquitin (PDB: 1UBQ) is a small protein (76 amino acids) that has received significant attention in the protein folding literature. We performed molecular dynamics simulations for this protein to use as a baseline, using a native-centric coarse-grained force field where every amino acid is represented by a single bead centered on the *α*-carbon; for more detail, see Appendix D. This formulation has been used to study protein folding in several previous studies *e.g*. (39; 35; 40). Our force field contains an adjustable scaling factor *η* which is determined by comparison to the experimental Gibbs free energy of folding. We ran replica exchange (REX) simulations and determined the ΔG of folding using the weighted-histogram analysis method (WHAM) (41), and found the parameter that reproduces the experimental Δ*G*_folding_ (−7.11 kcal/mol) taken from the literature (19).

We produced temperature quenching simulations using a Langevin integrator (friction parameter 0.05 ps^−1^, integration timestep 15 fs) and the OpenMM software package (42). We ran 200 independent trajectories consisting of 15 ns at 800 K, to induce temperature unfolding, followed by 300 ns at 298 K, to allow refolding. We printed trajectory snapshots every 5,000 timesteps (every 75 ps). The quenching trajectories were backmapped to a full backbone representation using PULCHRA (30), and analysed using the same procedure as the trajectories obtained from protein structure prediction methods.

## 3. RESULTS

### 3.1. Pathways from protein structure predictors are worse than chain length at predicting formal kinetics

We first evaluated whether the predicted pathways from protein structure prediction methods are consistent with experimental refolding kinetics. The methods were asked to classify if a protein chain folds through two-state kinetics or multistate kinetics; in other words, whether the folding reaction is fully concerted or progresses through an intermediate. The ground truth is a dataset of *in vitro* refolding experiments extracted from the literature.

As described in Methods, we modified the latest versions of seven state-of-the-art protein structure prediction methods to output their search trajectory. The first group, Rosetta and SAINT2, make use of a Monte Carlo minimization strategy based on fragment replacement. The second group, trRosetta, RaptorX, DMPfold and EVfold, use a flexible model with a simplified energy function as provided by CNS (43) or the Rosetta energy function (44), in combination with inter-residue restraints derived from co-evolutionary data. Of these, one model (EVfold) uses binary contacts predicted by a Potts model (45), while the other three use deep learning to predict inter-residue distances (DMPfold) and possible inter-residue orientations (trRosetta, RaptorX). The last method, RoseTTAFold, uses an iterative SE(3)-equivariant transformer that predicts protein structures in an end-to-end fashion without explicit minimization. These methods were used to produce 200 folding trajectories for each of the 170 proteins in our test set; except for the fragment replacement methods, SAINT2 and Rosetta, where due to high computational cost we generated only 10 trajectories per protein.

Generated pathways are influenced by the choices of the different protein structure prediction programs. Fragment replacement codes like SAINT2 and Rosetta start from the fully extended protein and slowly form compact states. Others like trRosetta and RaptorX start from a random conformation whose torsion angles have been selected from uniform sampling from a list of common torsion angles. RoseTTAFold initiates the trajectory in a compact structure that has been generated by inference on the MSA (and that often exhibits significant steric clashes). Despite the different initial states, all codes generate trajectories exhibiting complex folding dynamics.

The pathways were analysed using a method based on the fraction of native contacts between secondary structure elements. In a concerted, two-state mechanism, we expect a sudden change where most of the interactions between the secondary structure elements of a protein form in a single step, while in a multistate mechanisms we expect several sets of interactions forming at disjoint points of the trajectory. Our analysis (see Methods) identifies the steepest changes, and uses a statistical criterion to determine whether they should be considered as a single group (two-state) or multiple groups (multistate, where the interleading peaks can be regarded as intermediates). Table 1 shows the results of this classification.

**Table 1.** Performance of the different protein structure prediction methods at determining folding kinetics. Unsupervised metrics employ a simple rule *c*(**x**) that assigns a protein the most frequent kinetics *i.e*. if 50% or more of the decoys display multistate kinetics, the protein is taken to fold in multiple steps; otherwise it is considered two-state. Supervised metrics fit a logistic regression on *c*(**x**) and report the average of 1,000 5-fold cross-validation experiments; note that the supervised score may sometimes be worse than the unsupervised one if the model does not generalise well. The baseline is a logistic regression that uses only the length of the protein. Accuracy reports the average recall per class, to account for the slight imbalance of the dataset (90 two-state folders and 80 multistate folders). The F1-score is the harmonic mean of recall and precision. The area under the receiver-operating curve (AUROC) for length is computed by projecting the values to the [0,1] interval. We observe that chain length outperforms any of the protein structure prediction methods at predicting folding kinetics.

|  | RoseTTAFold | trRosetta | RaptorX | DMPfold | EVfold | SAINT2 | Rosetta | Length |
| --- | --- | --- | --- | --- | --- | --- | --- | --- |
| *10 decoys* |  |  |  |  |  |  |  |  |
| Unsupervised accuracy | 0.614 | 0.614 | 0.560 | 0.565 | 0.552 | 0.554 | 0.552 | - |
| Unsupervised F1-score | 0.637 | 0.588 | 0.472 | 0.679 | 0.525 | 0.586 | 0.513 | - |
| Supervised accuracy | 0.607 | 0.576 | 0.551 | 0.588 | 0.568 | 0.538 | 0.527 | **0.656** |
| Supervised F1-score | 0.637 | 0.620 | 0.558 | 0.667 | 0.643 | 0.620 | 0.655 | **0.731** |
| AUROC | 0.675 | 0.654 | 0.626 | 0.594 | 0.605 | 0.608 | 0.560 | **0.739** |
| *200 decoys* |  |  |  |  |  |  |  |  |
| Unsupervised accuracy | 0.623 | 0.546 | 0.576 | 0.556 | 0.608 | - | - | - |
| Unsupervised F1-score | 0.663 | 0.638 | 0.610 | 0.687 | 0.616 | - | - | - |
| Supervised accuracy | 0.612 | 0.573 | 0.563 | 0.581 | 0.610 | - | - | **0.656** |
| Supervised F1-score | 0.649 | 0.640 | 0.565 | 0.667 | 0.645 | - | - | **0.731** |
| AUROC | 0.669 | 0.631 | 0.602 | 0.622 | 0.658 | - | - | **0.739** |

Prediction accuracies are modest, but significant. Using a bootstrap test (*N* = 100, 000), we determined that all the structure predictors are significantly superior to a random classifier (AUROC = 0.500) at the 99% level of confidence. A randomised permutation test, however reveals that none of the predictors is significantly better at predicting folding kinetics than a linear classifier using only chain length. The fact that this sequence-agnostic classifier is better than any of the structure predictors suggests that, while protein structure prediction programs are capturing a non-trivial signal about folding, this signal is very weak.

The best predictor of folding kinetics appears to be RoseTTAFold (a deep learning model based on a transformer architecture which directly produces a structure from a multiple sequence alignment), closely followed by EVfold (based on energy minimization subject to evolutionary constraints). EVfold could be considered the most physically realistic method of those tested, since it does not modify the energy function to bias it towards the predicted native state. DMPfold is similar to EVfold, as it uses the same simulation engine (CNS), but the former uses a different method for introducing distance restraints: in DMPfold they are predicted with deep learning, whereas EVfold uses a Potts model. EVfold is a better predictor of folding kinetics than DMPfold, and also comparable to or better than RaptorX and trRosetta, which rely on deep learning. This suggests that, with the exception of RoseTTAFold, which belongs to a novel family of methods with physical assumptions baked into the model’s architecture, deep learning models are performing worse.

We also tested AlphaFold 2’s ability to predict folding kinetics, although in this case we had only one trajectory per protein. Using the method by Jumper *et al*. (14) we achieved an unsupervised accuracy of 0.613 and an unsupervised F1-score of 0.591 (note that other metrics, such as supervised cores or AUROC, since the score is binary due to the availability of only one trajectory per protein), which may hint at a similar performance to RoseTTAFold. If after averaging over multiple decoys the performance metrics remained constant then this would reinforce the notion that deep learning methods based on SE(3)-equivariance might be capturing folding information encoded in the multiple sequence alignment.

Overall the quality of the structure prediction output does not appear to relate to the ability of the method to classify folding kinetics (see Figure S2). In the 10 decoy dataset there is a tendency towards the methods that generate worse structure predictions also being worse at predicting kinetics, but this effect may be a product of reduced sampling. If we consider the 200 decoy dataset the method that has the lowest structure prediction accuracy, EVfold, is the second best predictor of kinetics. Similarly for a given program, the quality of the predictions is largely independent of model quality (see Figure S2).

We examined one of the methods that use deep learning, DMPfold, in more detail. DMPfold uses an iterative process where prior predictions are used to refine the potential employed in subsequent cycles. We compared the predictive power of multiple iterations, and observed that, while the area under the receiver-operating curve (AUROC) increases slightly with successive iterations, the overall accuracy is reduced (see Figure S3). The AUROC can be interpreted as the probability that a uniformly drawn two-state folder exhibits a higher proportion of two-state folding trajectories than a uniformly drawn multistate folder. This result suggests that, by iteratively refining predicted distances, the potential eliminates spurious predictions that might be a source of intermediates, as well as improve the final structure. However, since the accuracy is reduced, the description of the free energy hypersurface is not improved.

Finally, we found that some programs have an intrinsic bias towards predicting one or other folding mechanism. For example, for the majority of proteins, about 90% of the 200 DMPfold decoys exhibit two-state folding (hence the increase in AUROC from the 10 decoys sample to the 200 decoys sample), while RaptorX and EVfold tend towards predicting intermediates, and trRosetta presents a clear, but less marked bias towards two-state trajectories. These tendencies may explain the differences between unsupervised and supervised accuracy in Table S1.

Overall, these results suggest that protein structure prediction programs are not learning information about the folding mechanism.

### 3.2. Pathways from most protein structure predictors are uncorrelated with the rate constants of two-state folding

We next examined whether the protein structure prediction methods can predict the folding rate constant of the two-state processes. Our work follows that of Plaxco, Simons and Baker, who demonstrated that the average contact order of the native structure is strongly correlated with the folding rate constant of two-state proteins (46). Follow-up papers have suggested that other measures, such as fractions of secondary structure (47) or even predicted contacts (48), show similar correlations. We hypothesise that, if the folding pathways produced by protein structure methods were representative of folding, they should exhibit a similar relation, where the presence of the folding event in the trajectory is highly correlated with the folding rate constant.

We tested whether we could predict the folding rate constants of 79 two-state folding proteins from the PFDB (19) (see Appendix B for the experimental ground truth data). For each protein, we discarded all of the decoy trajectories that exhibited an intermediate and selected only two-state examples. In these trajectories, we localised the frame where the folding event started, and correlated its relative position in the full trajectory with the natural logarithm of the folding rate constant. As a baseline, we also computed the correlation with the average contact order and the chain length. We found that chain length outperformed average contact order at predicting the folding rate constant, counter to previous work that stated that length was not a useful predictor (46). This is potentially due to the use of different examples and increased dataset size (our dataset is six times the size of that in the original paper).

We found that most programs exhibit only a very weak correlation between the simulated trajectories and the folding rate constant. The Spearman correlation coefficients are not significant, at the 95% level of confidence, for trRosetta and RaptorX and DMPfold, and while EVfold, RaptorX and Rosetta display significant correlation, the correlation has the wrong sign: later folding events lead to larger (faster) rate constants. In contrast, the correlation between trajectories produced by RoseTTAFold and folding kinetics, although weaker in magnitude, has the correct sign. Nevertheless, all of the methods are significantly worse than the length of the protein chain at predicting the folding rate constant.

**Figure 3.**
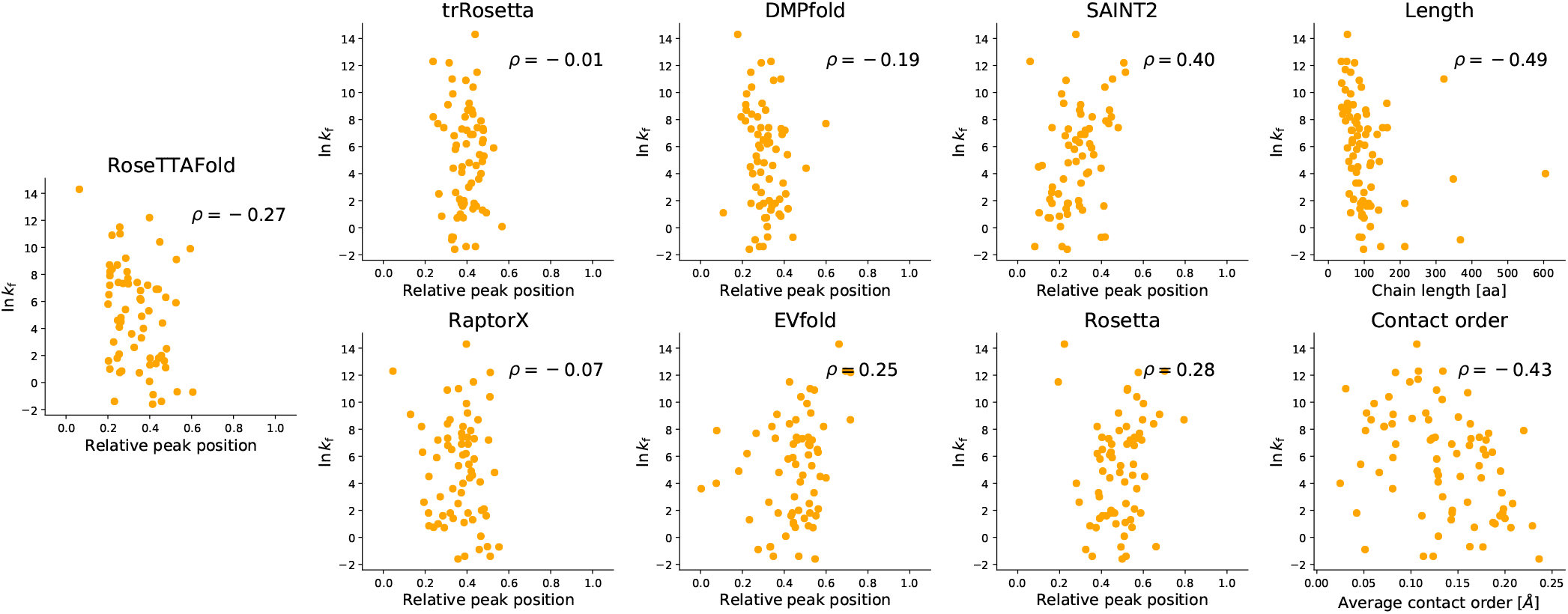
**Correlation between the folding rate constant** and folding events in simulated trajectories of the seven structure prediction methods considered, the length of the protein chain, and the average contact order of the native structure. Every point represents the average over the maximum number of decoys possible (200 decoys for RoseTTAFold, trRosetta, RaptorX, DMPfold and EVfold; and 10 decoys for SAINT2 and Rosetta).

We also found that AlphaFold 2 behaves similarly to RoseTTAFold, as found in the previous section. The Spearman correlation coefficient between the relative position of the folding event and the logarithm of the *k_f_* is −0.23, of the same order as RoseTTAFold and with the correct sign. Although the reduced number of decoys does not allow us to claim significance, the value suggests that the method is capturing some signal, and suggests that deep learning methods based on SE(3)-equivariance might detect the footprint that folding mechanisms have left in the multiple sequence alignment. However, it is unlikely that AlphaFold 2 would outperform the length of the protein chain at predicting the folding rate constant.

These results reinforce the conclusion that the ability of protein structure prediction methods to model folding pathways is inferior to trivial baselines.

### 3.3. Intermediates predicted by protein structure predictors are erratic and incompatible with available HDX data

As on occasion structure predictors do correctly identify folding kinetics we next examine if in these cases the structures predicted in the pathway are consistent with experimental data. We hypothesise that if the structure predictor has insight into the multistate process, it should (1) predict structures that are congruent with experimental measurements, and (2) produce consistent predictions of the intermediates across independent replicas for the same protein. Hydrogen-deuterium exchange (HDX) experiments probe unfolded regions of a protein at different stages of the folding process and allow us to identify which regions of an intermediate are structured and which have not yet folded (see Appendix C for details). We compared the predicted folding trajectories to these data.

We use the predicted trajectories to identify which pairs of secondary structure elements are interacting closely in the intermediate. This allows comparison between the noisy protein structure prediction pathways and the low structural resolution provided by experimental HDX data. For every protein and program, we consider a binary vector whose elements correspond to pairs of secondary structure elements that are in contact in the native structure. We then use the same trajectory analysis as in the previous section to identify which pairs interact in the folding intermediate (or, in the case of fructose-biphosphate aldolase A, the first intermediate). The metrics of these classifiers are summarised in Table 3.3.

**Table 2.** Performance of the structure predictors at identifying the secondary structure interactions present in an intermediate. The ground truth corresponds to a dataset of eleven proteins whose intermediates have been characterised with hydrogen-deuterium exchange (HDX) experiments. Accuracy reports the average recall per class, to account for the slight imbalance of the dataset. The Jaccard score reflects the average Jaccard similarity of the predictions, expressed as a binary string (where 1 means that the native contacts between secondary structure elements are formed in the intermediate, while 0 means they are not), with the true answer. The random baseline corresponds to an unbiased coin predicting whether two secondary structure elements are in contact.

|  | RoseTTAFold | trRosetta | RaptorX | DMPfold | EVfold | SAINT2 | Rosetta | Random |
| --- | --- | --- | --- | --- | --- | --- | --- | --- |
| *10 decoys* |  |  |  |  |  |  |  |  |
| Accuracy | 0.448 | **0.532** | 0.486 | 0.474 | **0.532** | 0.479 | 0.522 | 0.502 |
| F1-score | 0.210 | 0.192 | 0.217 | 0.000 | 0.125 | 0.000 | 0.039 | **0.252** |
| Jaccard | 0.052 | 0.052 | 0.052 | 0.054 | 0.052 | 0.052 | **0.166** | 0.094 |
| AUROC | 0.432 | **0.512** | 0.477 | 0.505 | 0.500 | 0.447 | 0.470 | 0.498 |
| *200 decoys* |  |  |  |  |  |  |  |  |
| Accuracy | 0.453 | 0.534 | 0.495 | 0.489 | **0.540** | - | - | 0.502 |
| F1-score | 0.222 | 0.169 | 0.110 | 0.026 | **0.307** | - | - | 0.252 |
| Jaccard | 0.052 | 0.052 | 0.052 | 0.052 | 0.052 | - | - | **0.094** |
| AUROC | 0.441 | 0.503 | 0.502 | 0.492 | **0.530** | - | - | 0.498 |

Intermediate structures are predicted with very low accuracy by all methods. A randomised permutation test shows that only one of the predictors, EVfold, exhibits predictive power superior to the random baseline. In contrast, RoseTTAFold is significantly worse than the random sample. This suggests that deep learning models are not learning the physics of folding, but rather collecting statistical information about crystal structures.

As an additional sanity check, we considered whether the structures generated throughout the trajectories are consistent with basic physical rules. We computed the clashscore (49) of every snapshot in the first ten decoys using Phenix (50) and compared them against a threshold value of 30 clashes per 1,000 atoms, determined as the 99th percentil of PDB structures with resolution ≤ 2.5*Å* (see Figure S5). We observed that the majority of the methods produce a large number of structures with large clashes: methods based in CNS like DMPfold and EVfold produced over 80% of unphysical structures, and even the best methods like RaptorX and AlphaFold produced nearly 30-40% of structures with clashing atoms. This finding suggests that the potentials generated are not considering basic physical principles throughout the intermediate stages of the predictive process. This may explain the relative bad quality of intermediate predictions with respect to predictions of formal kinetics or the folding rate constant.

We then examined the variation between the predicted interactions by computing the Jaccard similarity between the binary vector of predicted interactions and the ground truth. This similarity is very low, in most cases worse than random, suggesting that independent replicas of the folding pathway by the protein structure prediction methods often lead to markedly different structural intermediates. These results once again imply that while the predictors may be good at modeling the energy hypersurface around the global minimum, they are not capturing other attractors and therefore produce erratic pathways.

The comparison with AlphaFold 2 suggests that the latter produces similar results. Of the nine proteins, seven are predicted with a Jaccard similarity of ~ 0.1 to the ground truth (see Figure S6). The two proteins that are predicted with some accuracy, horse cytochrome C and cardiotoxin analogue III, are also the smallest in the dataset, which once again raises a concern of reduced entropic pressure. This suggests that AlphaFold 2 does not present any advantage at predicting the folding intermediates of a protein chain.

We then investigated if these results extend from the proteins with HDX annotations, to the entire dataset of proteins we simulated. We computed the binary vectors for all pathways of multistate proteins exhibiting an intermediate, and computed the average Jaccard similarity for every protein (Figure 4). The average pairwise Jaccard similarity is 0.1, and in most cases there are only a handful of proteins with an average over 0.5. The yeast cell-cycle control protein p13suc1 (PDB: 1PUC) is one of this handful; it presents only four native interactions, suggesting that this is again due to reduced entropic pressure. Overall, the pathways produced by protein structure prediction methods are erratic and generally inconsistent, suggesting that any ability to correctly predict multistate behaviour does not arise from an understanding of the intermediates in the folding pathway.

**Figure 4.**
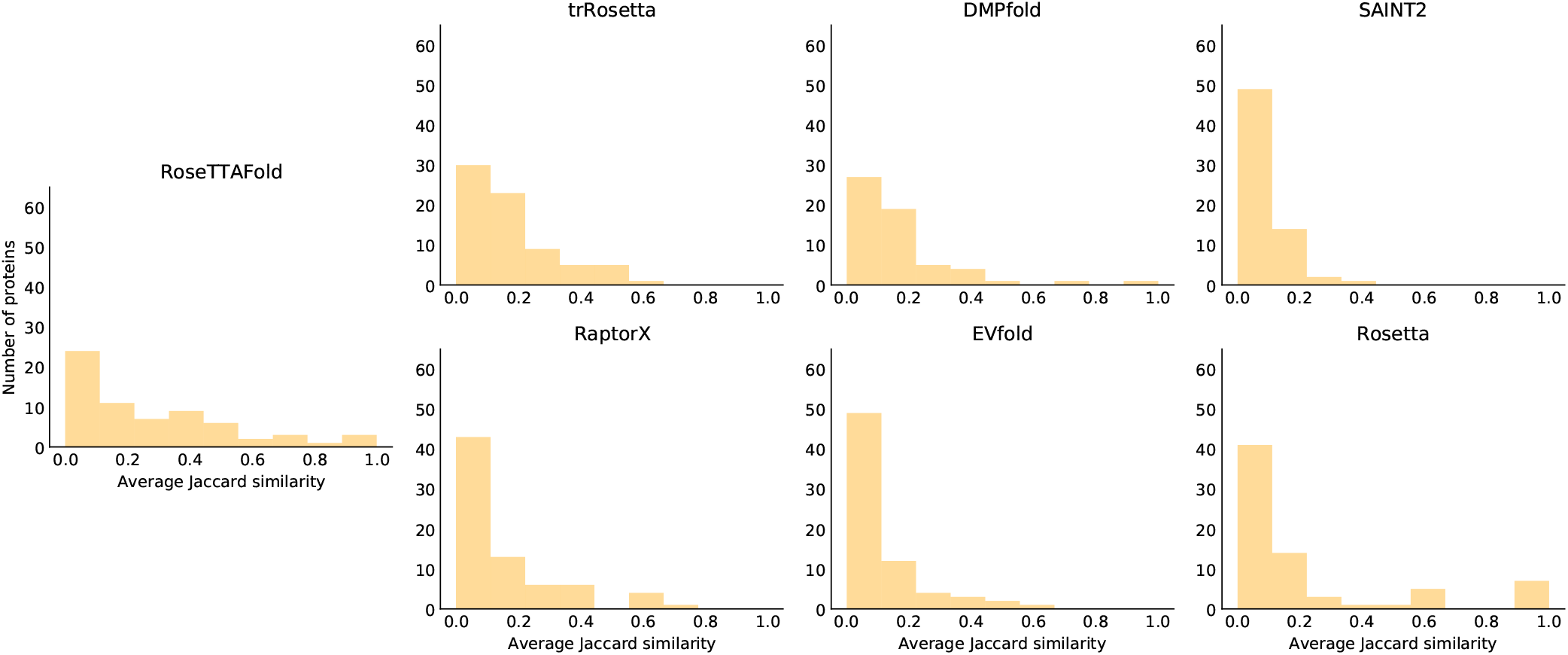
Average pairwise Jaccard similarity between multistate folding trajectories across all proteins in the dataset, for the seven structure prediction programs. Most methods exhibit significant variability between independent trajectories.

### 3.4. Case study: ubiquitin

Ubiquitin is a 76-residue protein involved in signaling, notably by marking proteins for degradation by the 26S proteasome (51). The folding and dynamics of ubiquitin have been widely studied both experimentally (21) and computationally (52; 53), including by millisecond all-atom unbiased molecular dynamics simulations of folding (54). The kinetics of folding were an object of controversy in the late nineties, with claims of a three-state mechanism (55), although the consensus opinion in the literature is now that ubiquitin is a two-state folder (56; 57).

We analysed the protein structure prediction trajectories for ubiquitin generated during our analysis, and as a physically-inspired baseline we considered a coarse-grained molecular dynamics (CGMD) simulation. Representative trajectories for each program are provided in Supplementary Videos 1 to 9. We found that ubiquitin displays two-state folding kinetics in most of the simulated trajectories of RoseTTAFold, trRosetta, DMPfold and EVfold, but only in less than one quarter in RaptorX (see Figure 5b). Surprisingly, CGMD simulations also suggested that the folding is multistate, with 62% of the trajectories exhibiting an intermediate. These results suggest that all codes, except potentially RoseTTAFold and DMPfold, are generating stable intermediates that do not reflect experimental kinetics.

**Figure 5.**
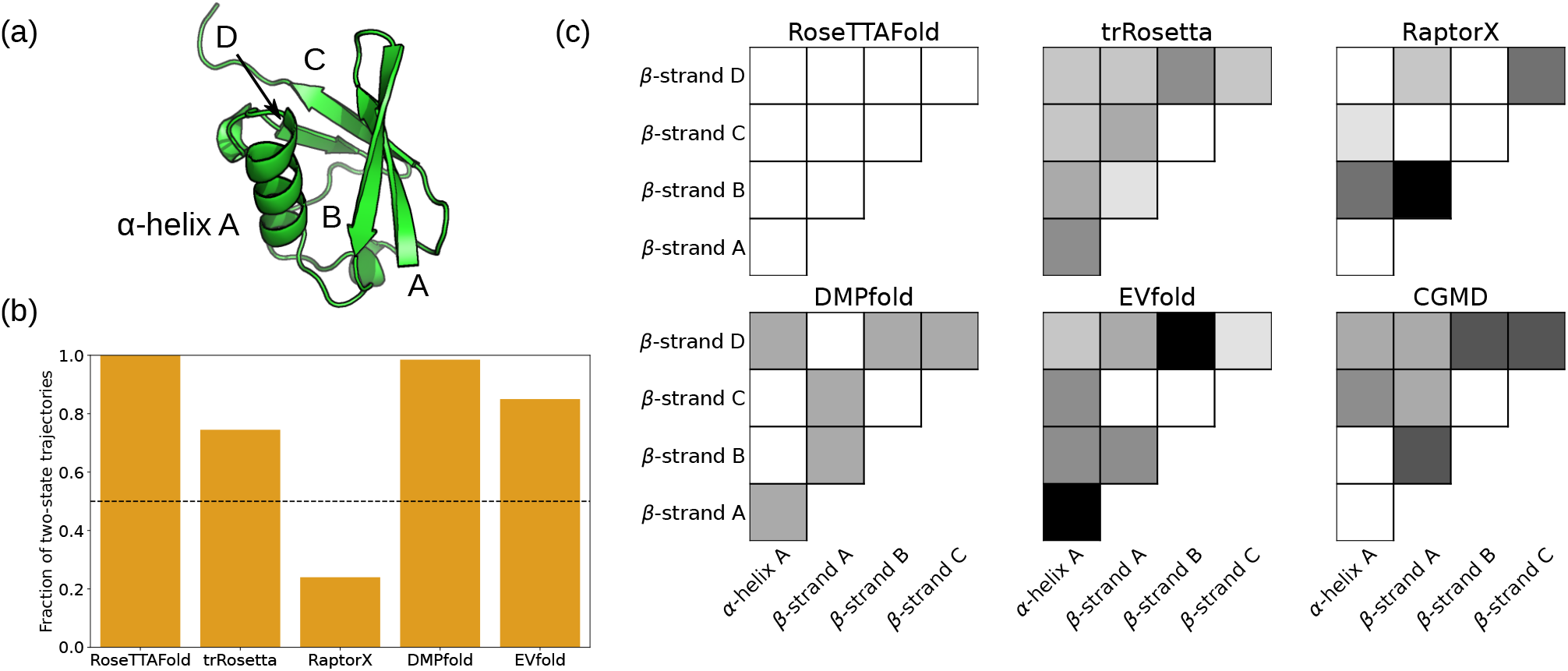
Simulations of human ubiquitin folding. (a) Native structure of human ubiquitin (PDB: 1UBQ). (b) Proportion of trajectories generated by each program that exhibit two-state dynamics. (c) Pairs of secondary structure elements that are formed in the identified intermediate. Color represents the proportion of trajectories where an intermediate presents a given interaction, where white is 0 (does not appear in any trajectory) and black is 1 (appears in all trajectories). Note that none of the RoseTTAFold trajectories exhibits an intermediate, hence none of the interactions is identified.

We then examined the structures of the intermediates, and found significant variability. Most programs, including the CGMD reference, have a tendency towards the *β*-strands A and B interacting (see Figure 6a), signaling the formation of a *β*-hairpin. This feature has been observed experimentally in NMR studies of unfolded ubiquitin (58) in up to 8M urea, and could suggest that the predictors are identifying strong interactions that govern the formation of metastable structures. However, most of the structure predictors also exhibit some spurious interactions that are not observed in either the CGMD reference or the experimental data, such as interactions between the *α*-helix and the *β*-strands A and B (see Figure 6a). This suggests that, while the structure predictors do capture some interesting interactions, these are likely to be of limited use, since they are hidden within large amounts of spurious information.

**Figure 6.**
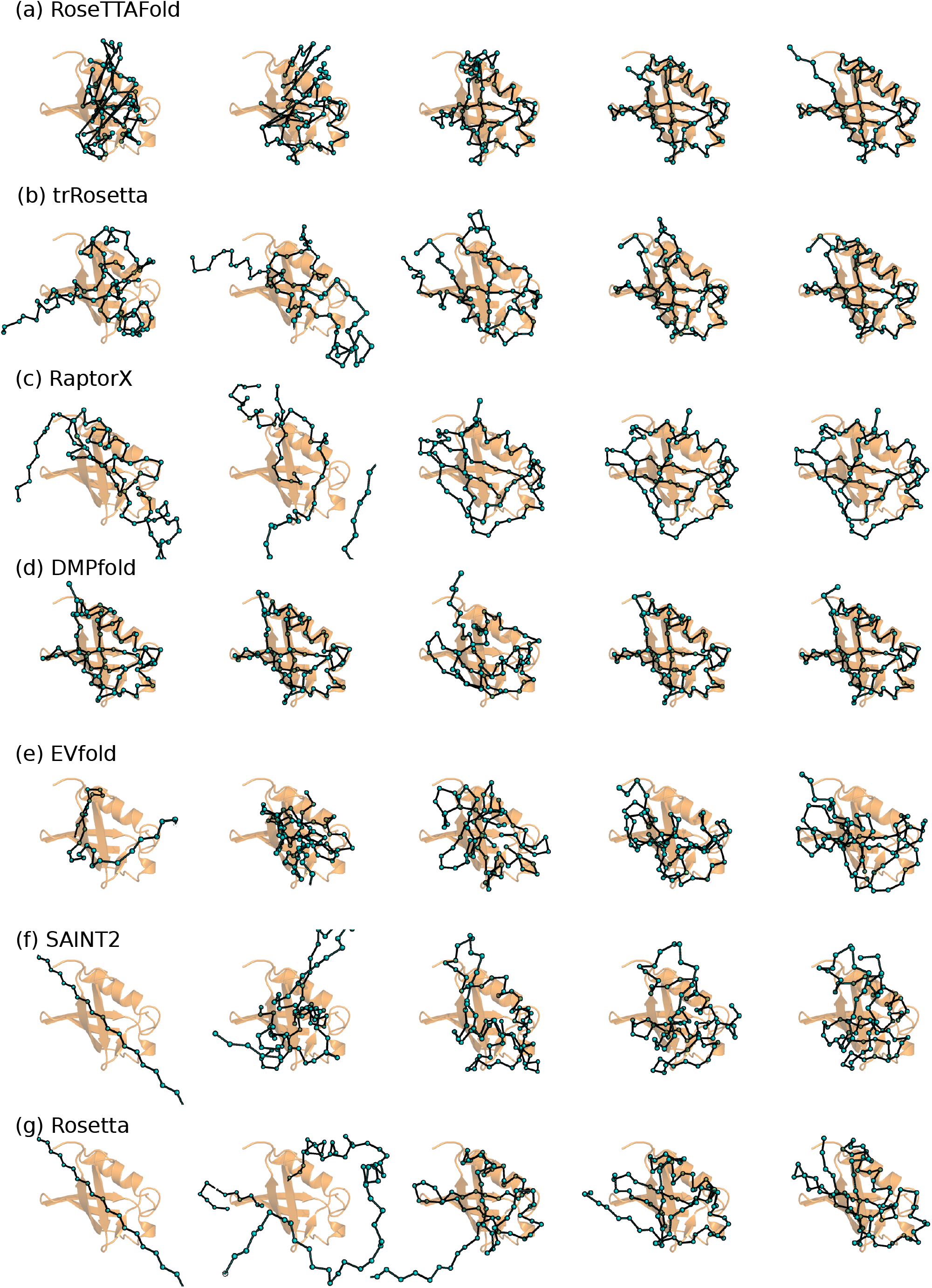
Representative snapshots of the simulated folding trajectories for human ubiquitin, for the seven structure prediction programs. The *α*-carbons are represented as blue beads joined by dark rods. Every structure has been aligned to the crystal structure (PDB: 1UBQ), which is shown as an orange cartoon in the background. are inconsistent with all available experimental data, in terms of folding mechanism, kinetics or structural data. In the general context of computational protein biophysics, our results suggests that current protein structure prediction programs, while now very successful at their primary role, are not an appropriate tool to investigate folding.

We next inspected the simulated trajectories visually. As Figure 6 shows, despite RoseTTAFold exhibiting clear two-state dynamics according to the collective variables, all of its trajectories display unphysical behaviour. Initial structures are characterised by unrealistic steric clashes and violations of the geometry of the peptide bond, and are followed by a relaxation of the backbone into the final structure.

trRosetta and RaptorX present trajectories with comparable dynamics, probably due to the similarities between their protocols. In these trajectories, the protein exhibits random movements in the unfolded state, followed by collapse and formation of secondary structure, concomitant with the activation of the biasing potential, and finally small local exploration of the positions of the loops. This mechanism is consistent with the CGMD simulations, although in several snapshots the protein gets trapped in local minima of the biasing potential, which gives rise to detected intermediates.

DMPfold and EVfold, despite using a similar approach (the CNS optimisation engine in combination with predicted contacts), exhibit very different folding trajectories. DMPfold starts in a structure consistent with predicted distances, and explores the neighbourhood of this structure, whereas EVfold starts in an elongated state, experiences collapse, and then explores several potential conformations, with no apparent relation between them, until it finds the best structure according to its energy function. Both mechanisms are inconsistent with the CGMD simulations.

The AlphaFold 2 trajectory is similar to RoseTTAFold. The initial frames exhibit significant steric clashes, which are resolved after about 10 Evoformer iterations, and are followed by small oscillations around the equilibrium structure that are reminiscent of a protein subject to harmonic restraints. These results once again support the hypothesis that these methods are not learning the free energy hypersurface, but only the small free energy funnel that surrounds the native state.

## 4. DISCUSSION

In this manuscript we have investigated whether state-of-the-art protein structure prediction methods can provide any insight into protein folding pathways. We generated tens of thousands of folding trajectories with seven protein structure prediction programs (RoseTTAFold, trRosetta, RaptorX, DMPfold, EVfold, SAINT2 and Rosetta) and obtained a set of AlphaFold 2 trajectories, and used them to determine major features of folding using a simple set of statistical rules. We found that protein structure prediction methods can in some cases distinguish the folding kinetics (two-state vs multistate) of a chain better than a random baseline, but not significantly better, and often significantly worse, than a simple, sequence-agnostic linear classifier using only the number of amino acids in the chain.

Using a similar approach, we examined the relationship between simulated trajectories and other experimental observables: the folding rate constant of two-state folders, and the structure of intermediates in multistate trajectories. The simulated trajectories were in most cases not better than random at predicting the contacts formed in an intermediate, and in the case of predicting folding rate constants, none of the methods was superior to a linear classifier using the length of the protein chain.

Our results demonstrate that state-of-the-art protein structure prediction methods do not provide an enhanced understanding of the principles underpinning folding. Simulated trajectories from protein structure prediction methods There are some limitations to our study. First of all, the concepts of folding intermediate and folding formal kinetics are imprecise. For example, many proteins have a tendency to form compact, molten globule structures, that may then fold cooperatively in a process that is referred to as “two-state” (*e.g*. 59). The folding mechanisms of multiple proteins have been widely discussed in the literature with conflicting results, (*e.g*. for ubiquitin (21) or T4 lysozyme (60; 61; 62)). Folding is itself highly sensitive to an array of experimental conditions that includes temperature, pH and concentration of denaturant, and it may be difficult to discern when the methods are not correctly modeling the physics or simply portraying the wrong conditions.

While our results have shown the lack of consistency between the folding trajectories generated by protein structure prediction methods and experimental data, we have also seen that most structure predictors are better than random suggesting that a weak signal exists. The next stage will be to investigate how to extract the limited amount of folding information that is encoded in current protein structure prediction programs.

## ACKNOWLEDGEMENTS

The authors would like to thank the AlphaFold 2 team at DeepMind for providing folding trajectories for analysis. C. O. would also like to thank Oliver Crook for advice on the statistical analysis of significance, and F. Hoffmann-La Roche, UCB and EPSRC (EP/M013243/1) for financial support.

## APPENDIX

## A. EXPERIMENTAL FOLDING KINETIC DATA

**Table S1.**
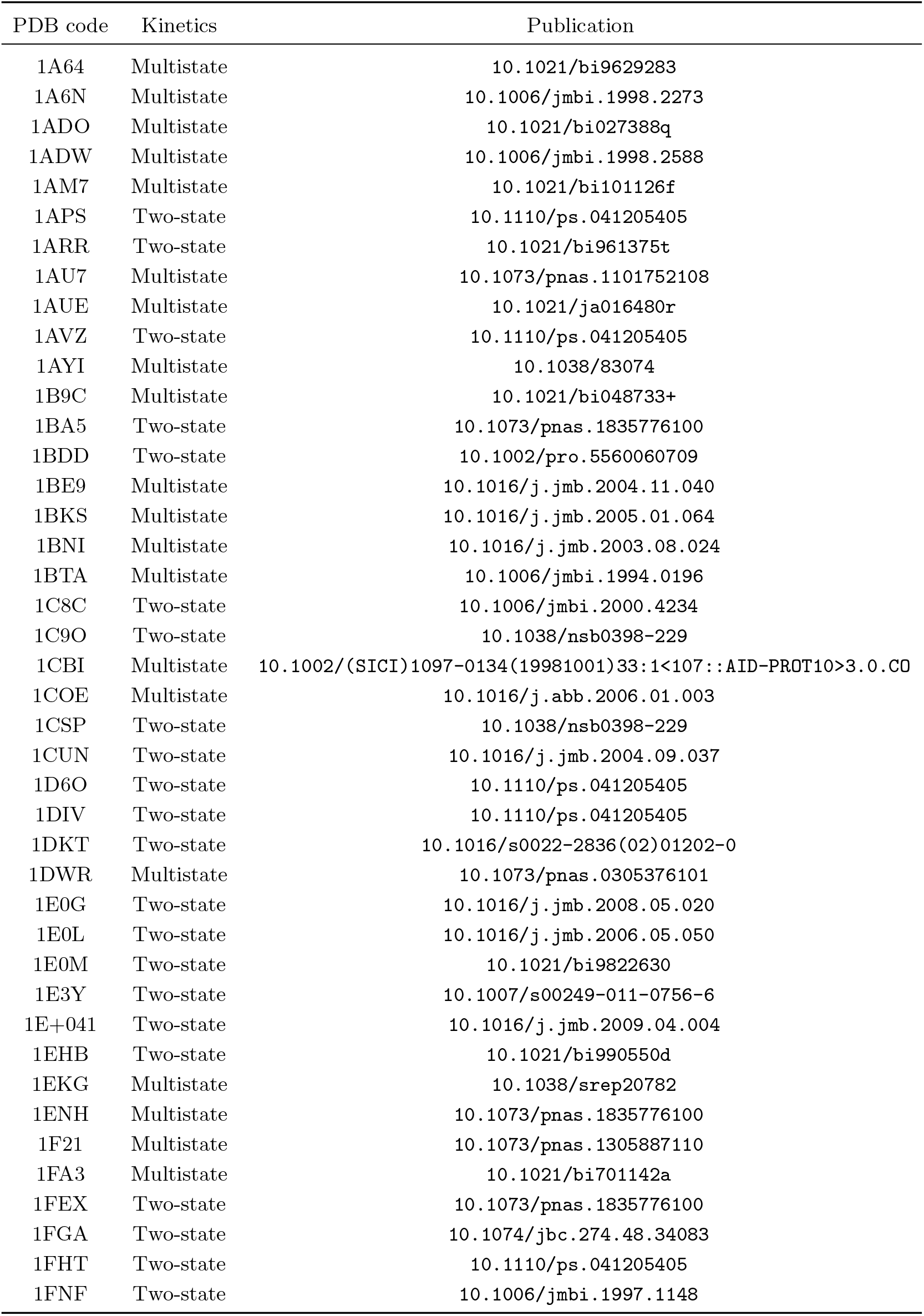
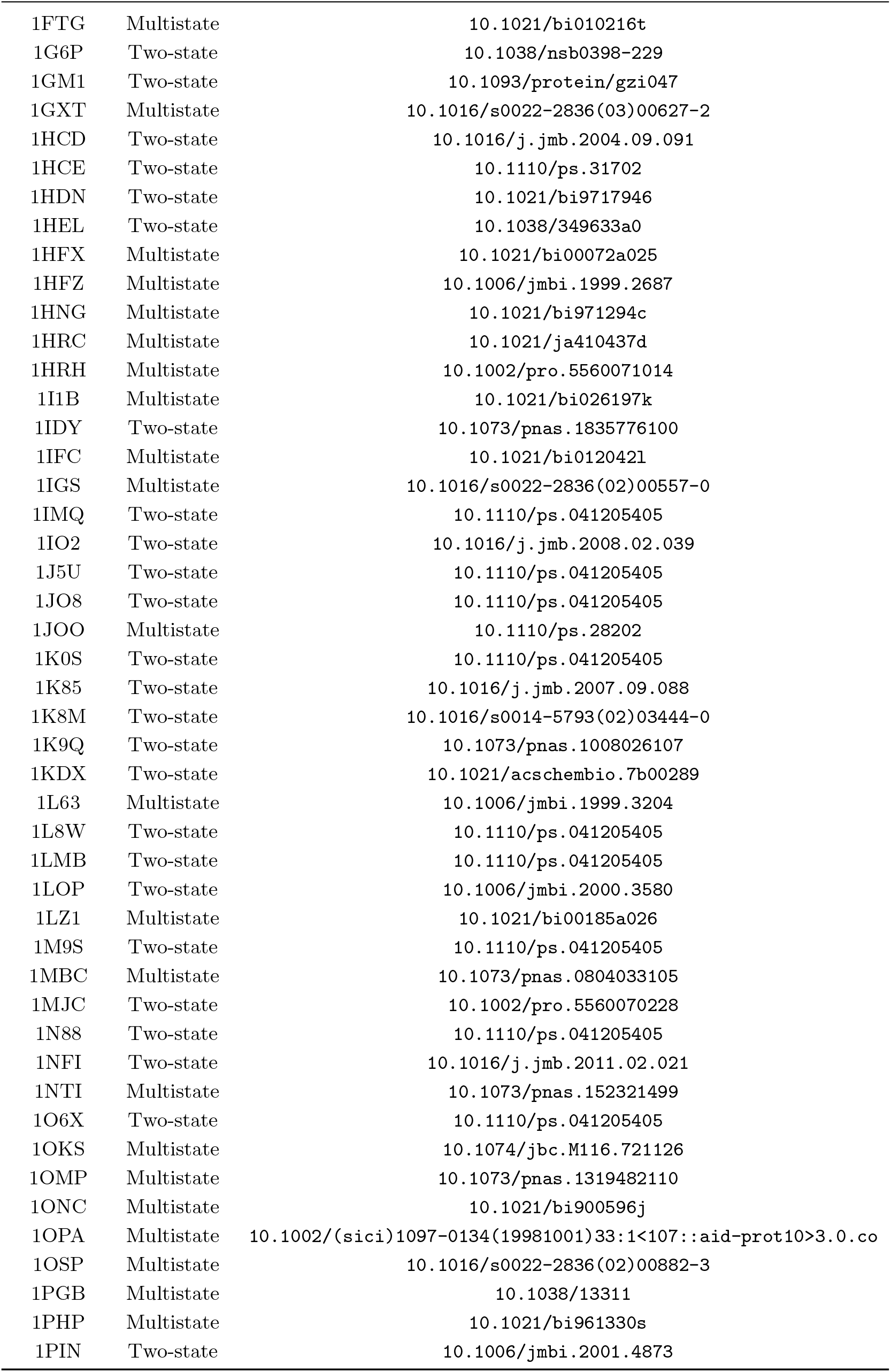
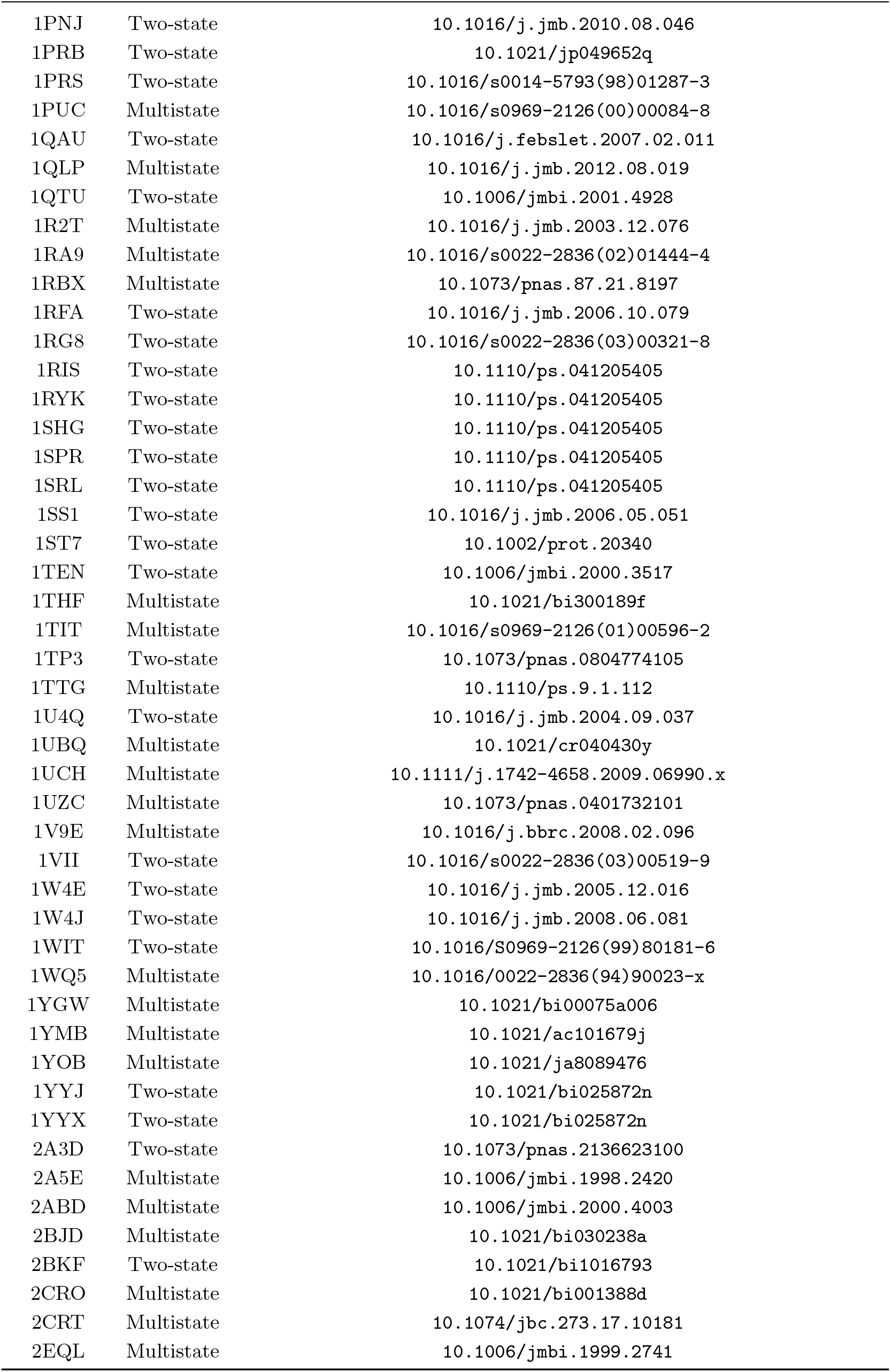
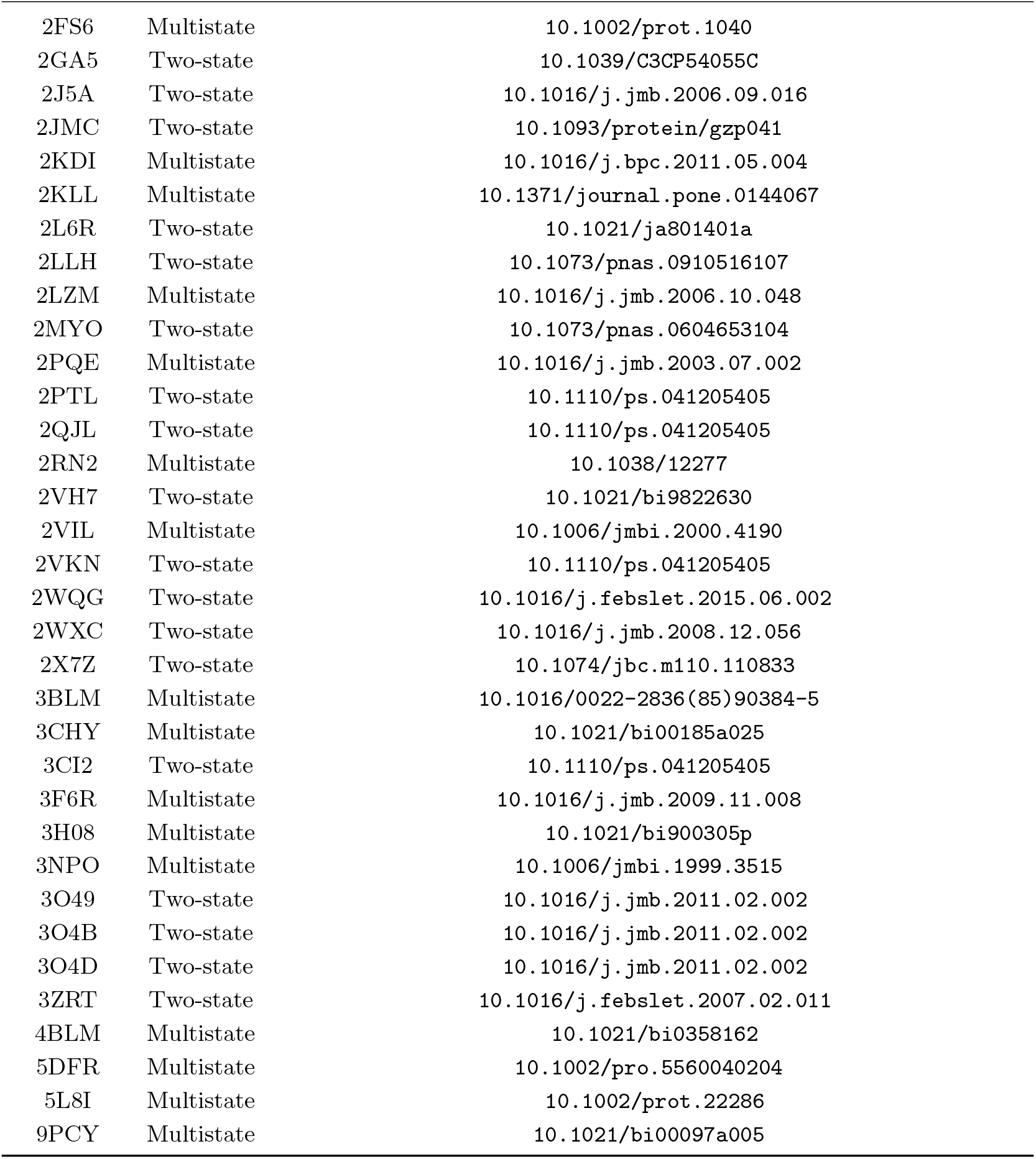
Experimental folding data used in this work

## B. EXPERIMENTAL TWO-STATE FOLDING RATE CONSTANTS

**Table S2.**
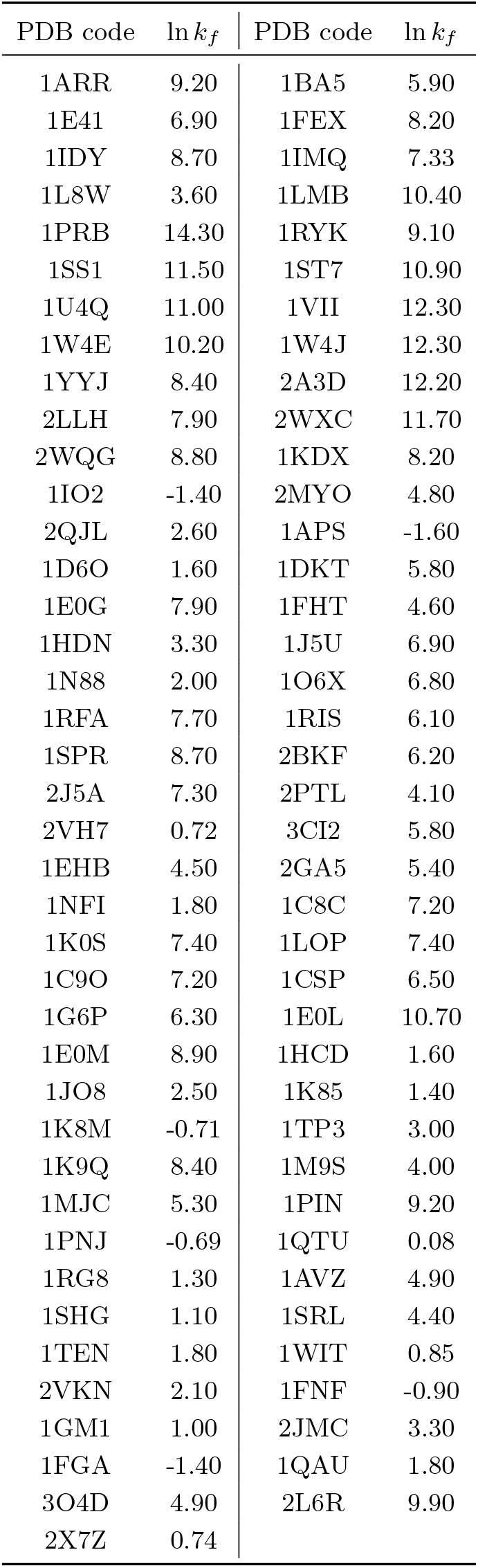
Folding rate constants

## C. EXPERIMENTAL STRUCTURAL FOLDING DATA

### C.1. Fructose-bisphosphate aldolase A

Pan and Smith (63) studied the folding of rabbit muscle aldolase (PDB: 1ADO) using HDX-MS. The authors proposed a folding mechanism whereby an initial collapsed state is formed cooperatively from the union of four widely separated regions of the backbone, followed by two sequential folding steps of individual domains. The authors annotated the regions of the protein as corresponding to each of these intermediates, by means of the peptides obtaining during HPLC-MS analysis.

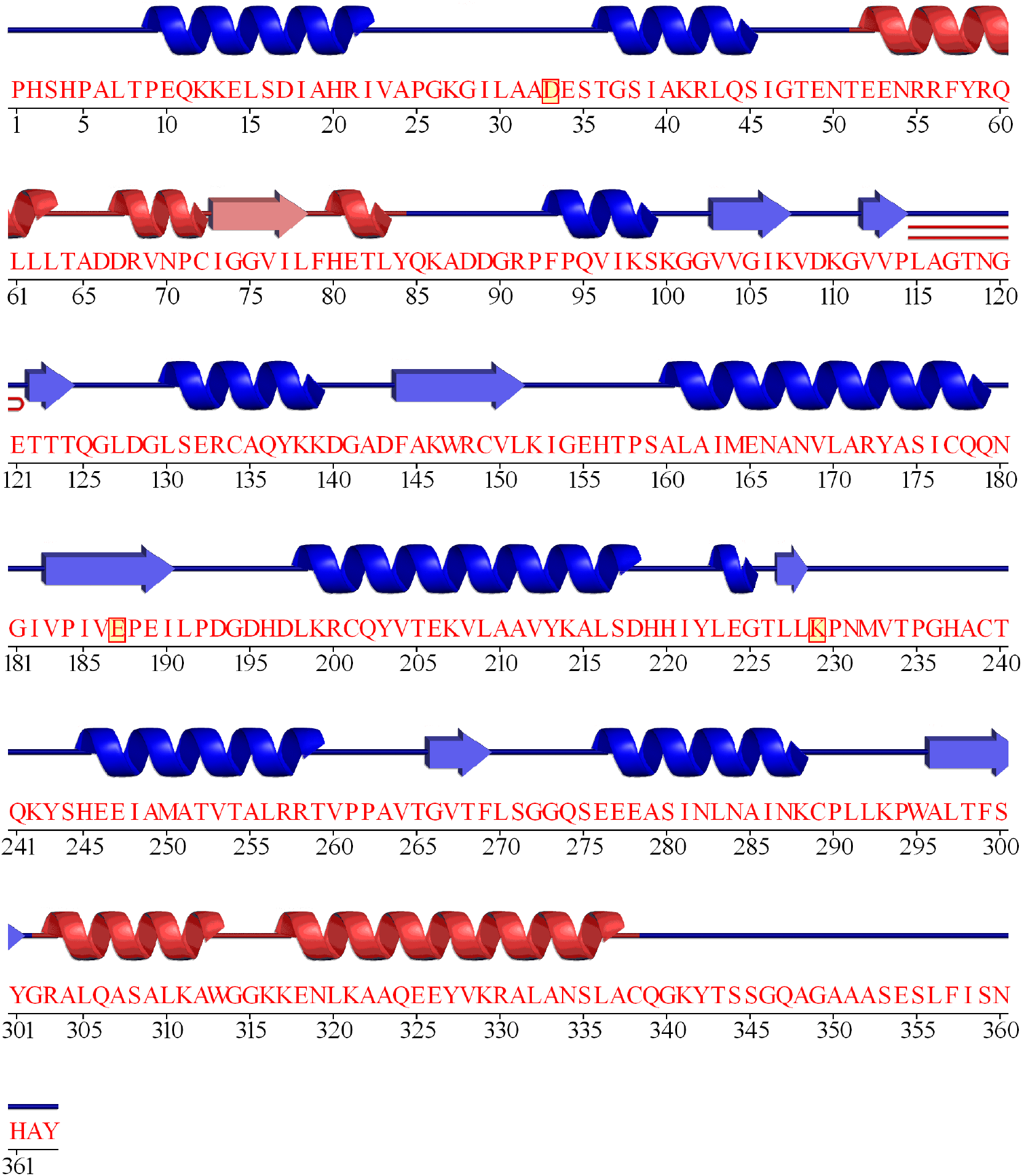

### C.2. Alpha-subunit of tryptophan synthase

Wintrode et al. (64) studied the folding of the *α*-subunit of tryptophan synthase (PDB: 1BKS, for an E. coli homologous with 85% sequence identity), a TIM barrel protein from *E. coli*, using HDX-MS. The authors identify an obligate intermediate, which comprises most of the N-terminal region.

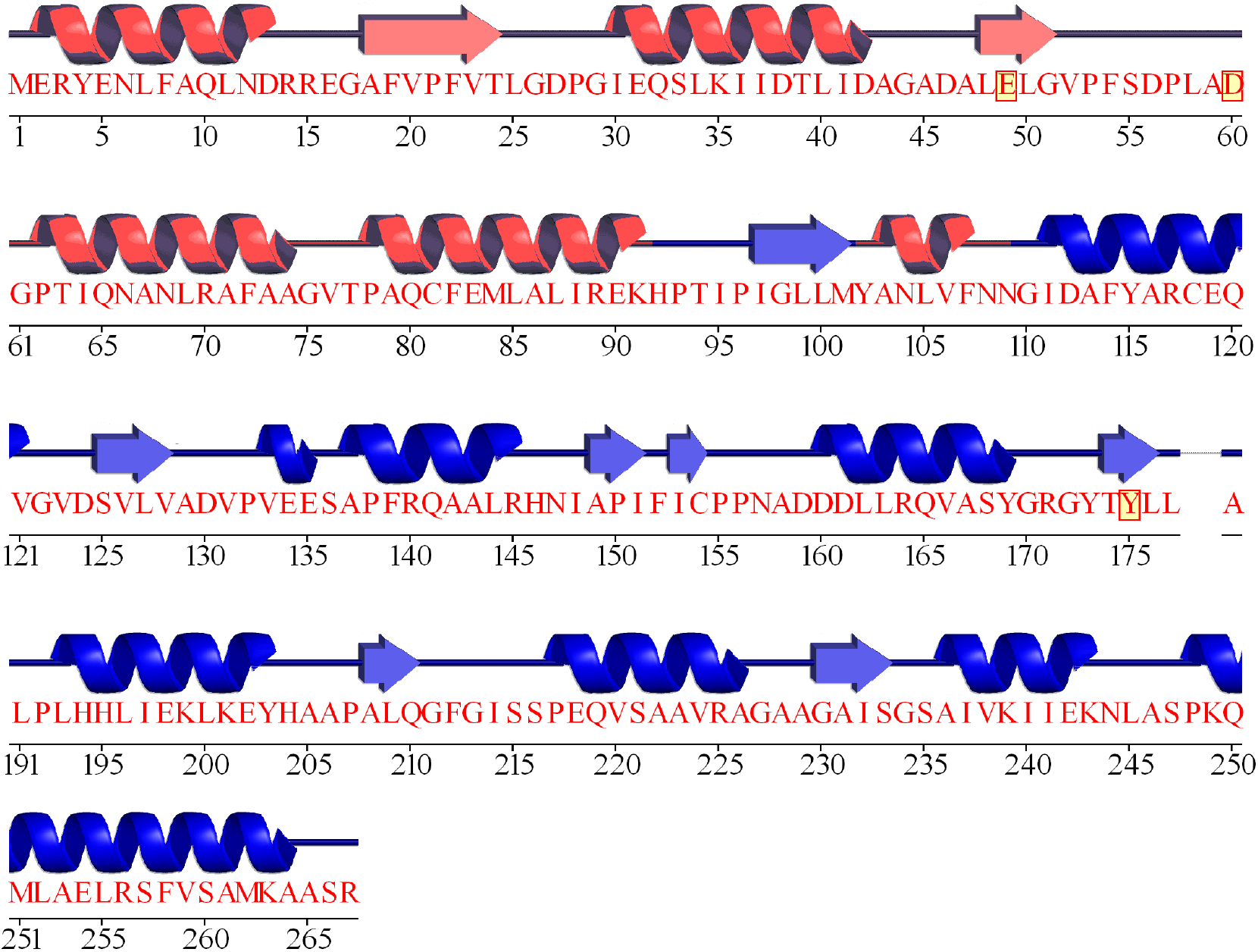

### C.3. Cytochrome C

Roder et al. (65) studied the folding of horse’s cytochrome C (PDB: 1HRC) using HDX-NMR. The authors observe that the contact between the C-terminal and N-terminal helices is formed early in folding, followed by structuring of the rest of the protein. A study by Elove et al. (66) identified multiple possible folding pathways originating from different coordinations to the heme group. A study by Fazelinia et al. (67) investigated folding during the first 140 *μ*s using a microfluidics device and HDX-NMR, showing that interactions between *α*-helices drives condensation at the start of folding.

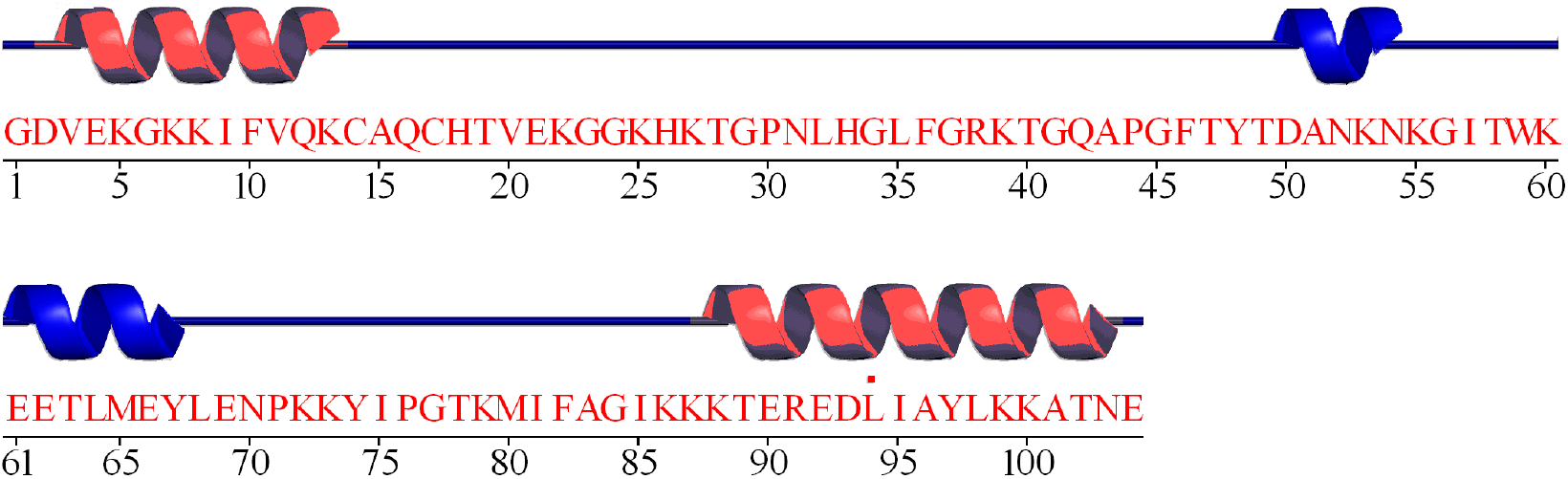

### C.4. Staphilococcal nuclease H124L

Walkenhorst et al. (68) studied the folding of staphilococcal nuclease with a H124L mutation (PDB: 1JOO). Their data shows that the formation of the *β*-barrel domain, in particular the *β*-hairpin formed by strands 2 and 3 and a site in the C-terminus, precedes the formation of the *α*-helical domain.

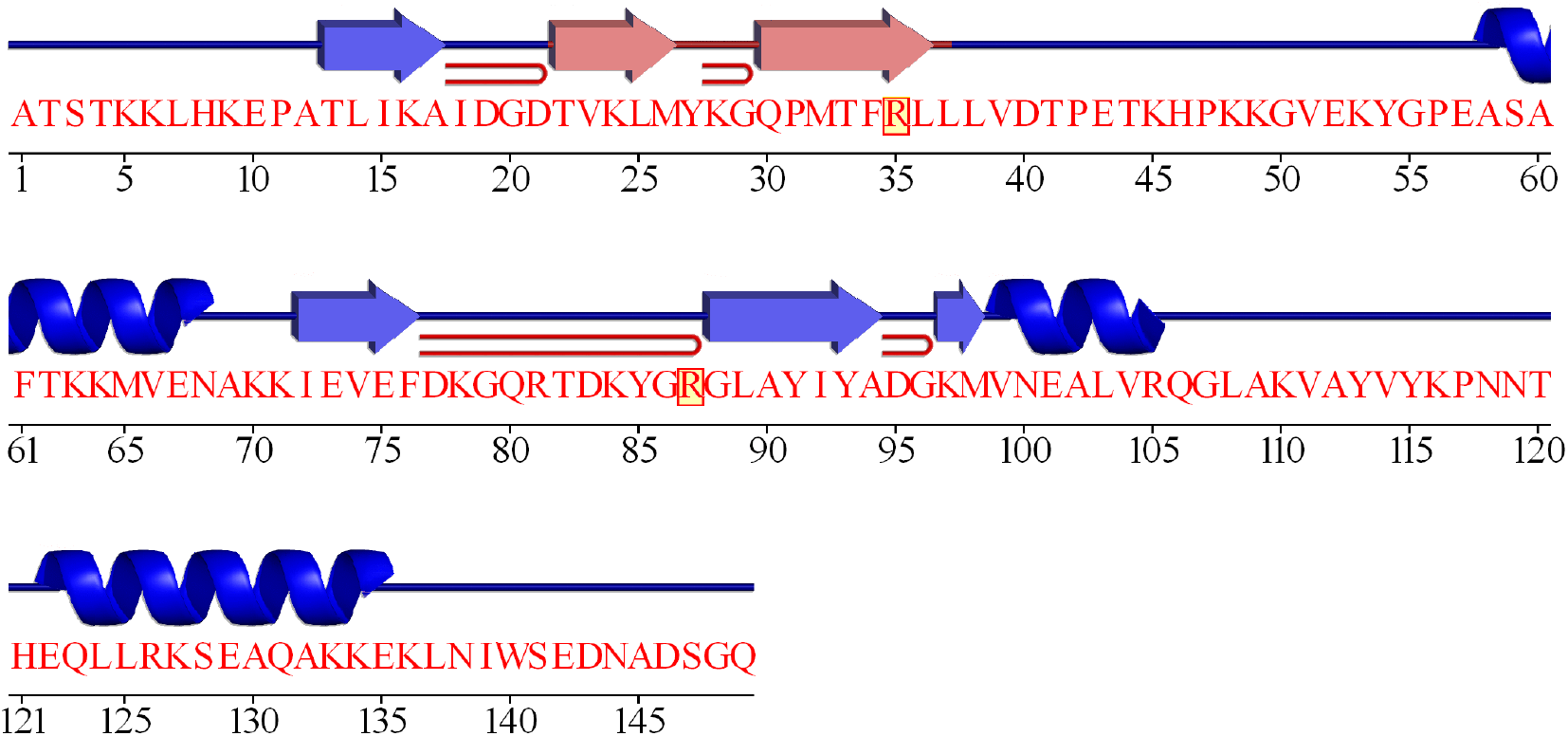

### C.5. Triosephosphate isomerase

Pan et al. (69) studied the folding of rabbit triosephosphate isomerase (PDB: 1R2T) using HDX-MS. They found that the C-terminal half folds to form the intermediate, which then forms a TIM barrel with the N-terminal half.

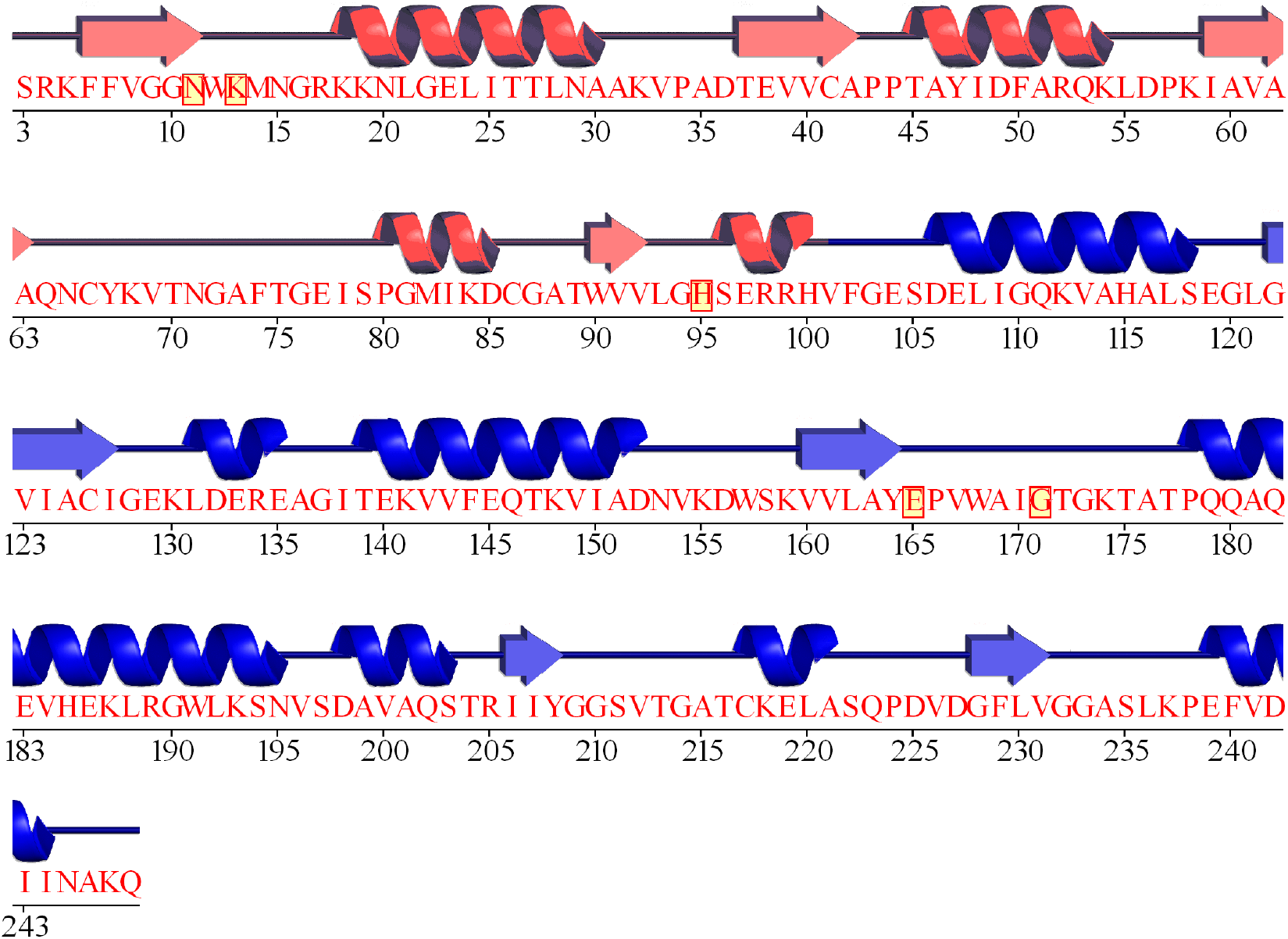

### C.6. Ribonuclease A

Udgaonkar et al. (70) analysed the folding of bovine pancreatic ribonuclease A (PDB: 1RBX) using HDX-NMR. They identify an intermediate with the first N-terminal *α*-helices unformed.

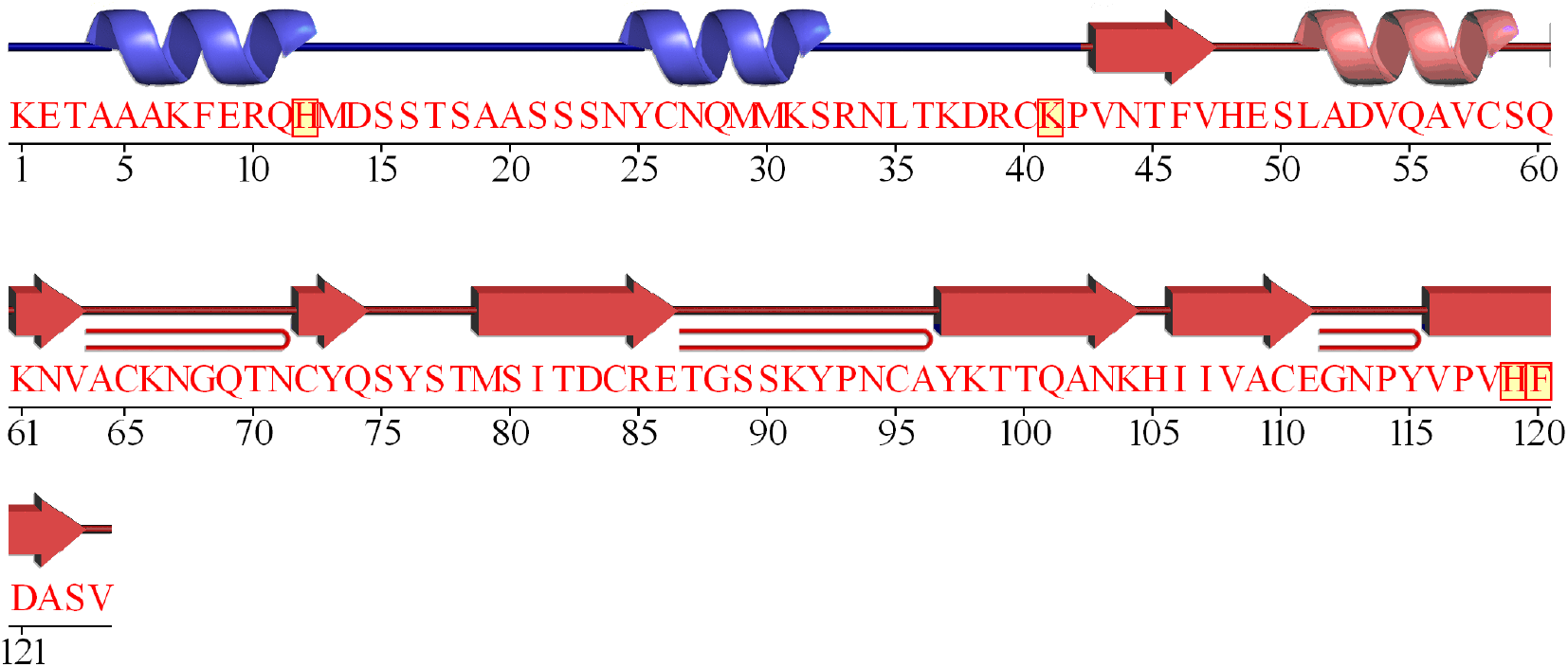

### C.7. Myoglobin

Pan et al. (71) studied the folding of horse apo-myoglobin (PDB: 1YMB) using HDX-MS.

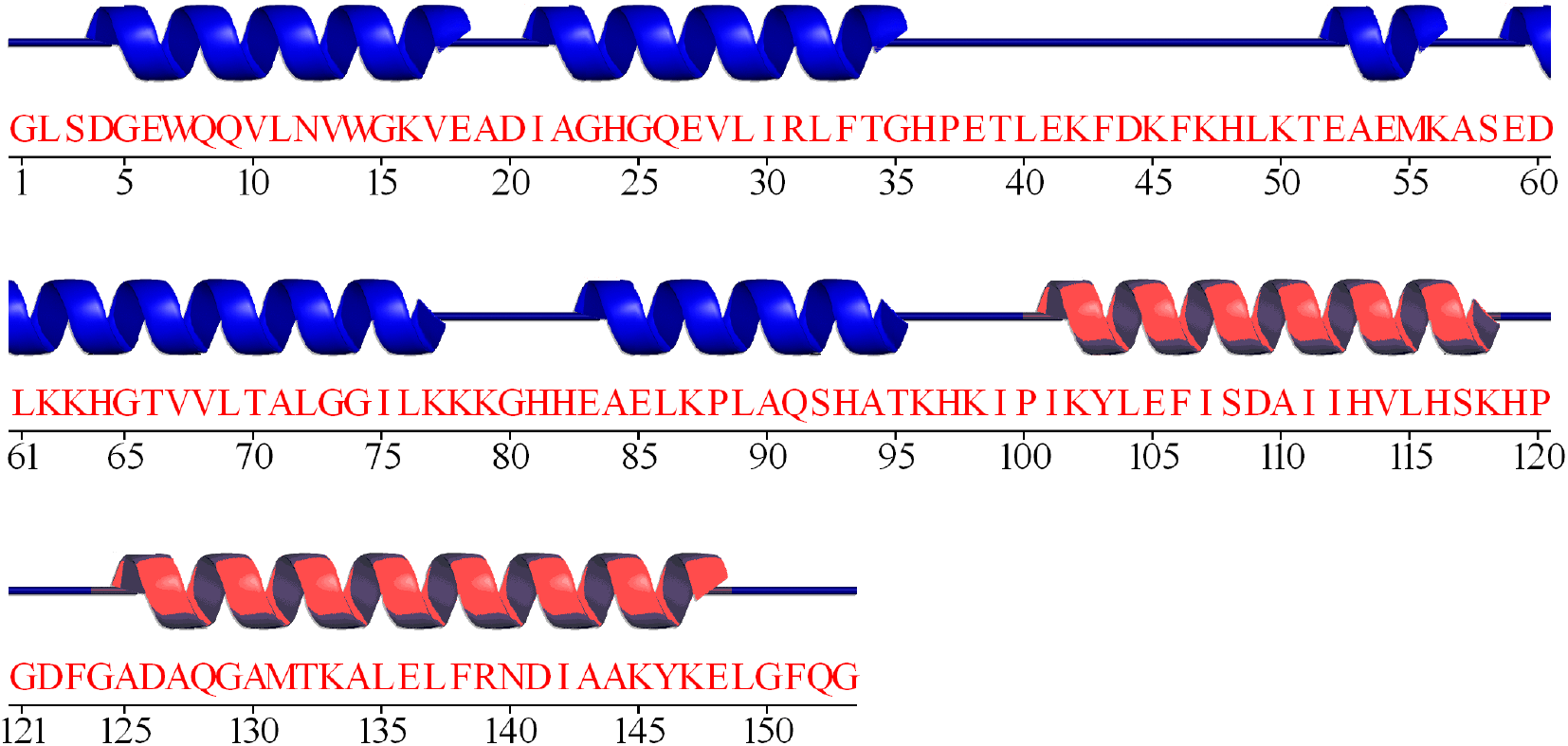

### C.8. Cardiotoxin III

Sivaraman et al. (72) studied the folding of cardiotoxin analogue III (PDB: 2CRT), a protein present in the venom of *Naja naja atra*. This protein is a small all-*β* protein with two recognisable *β*-sheets: a double-stranded *β*-sheet closer to the C-terminus, and a triple-stranded *β*-strand closer to the N-terminus. Experimental results show that the triple-stranded *β*-sheet folds about 10 ms faster than the double-stranded element.

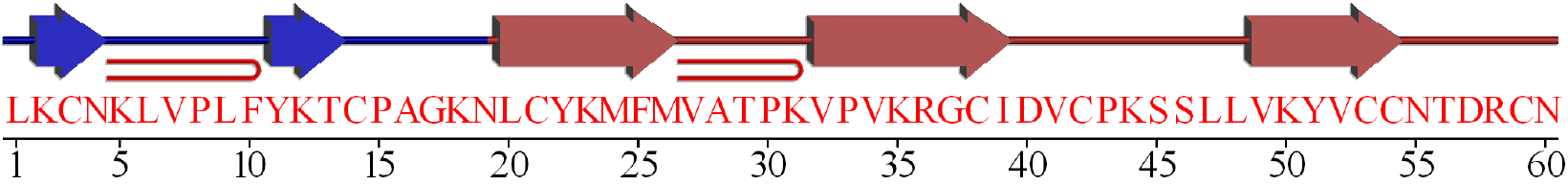

### C.9. Flavodoxin-2

Nabuurs et al. (73) studied the folding of flavodoxin 2 (PDB: 1YOB) from *A. vinelandii* using HDX-NMR. The authors identify an off-pathway intermediate where most of the secondary structure is formed, except for two regions that adopt a random coil conformation.

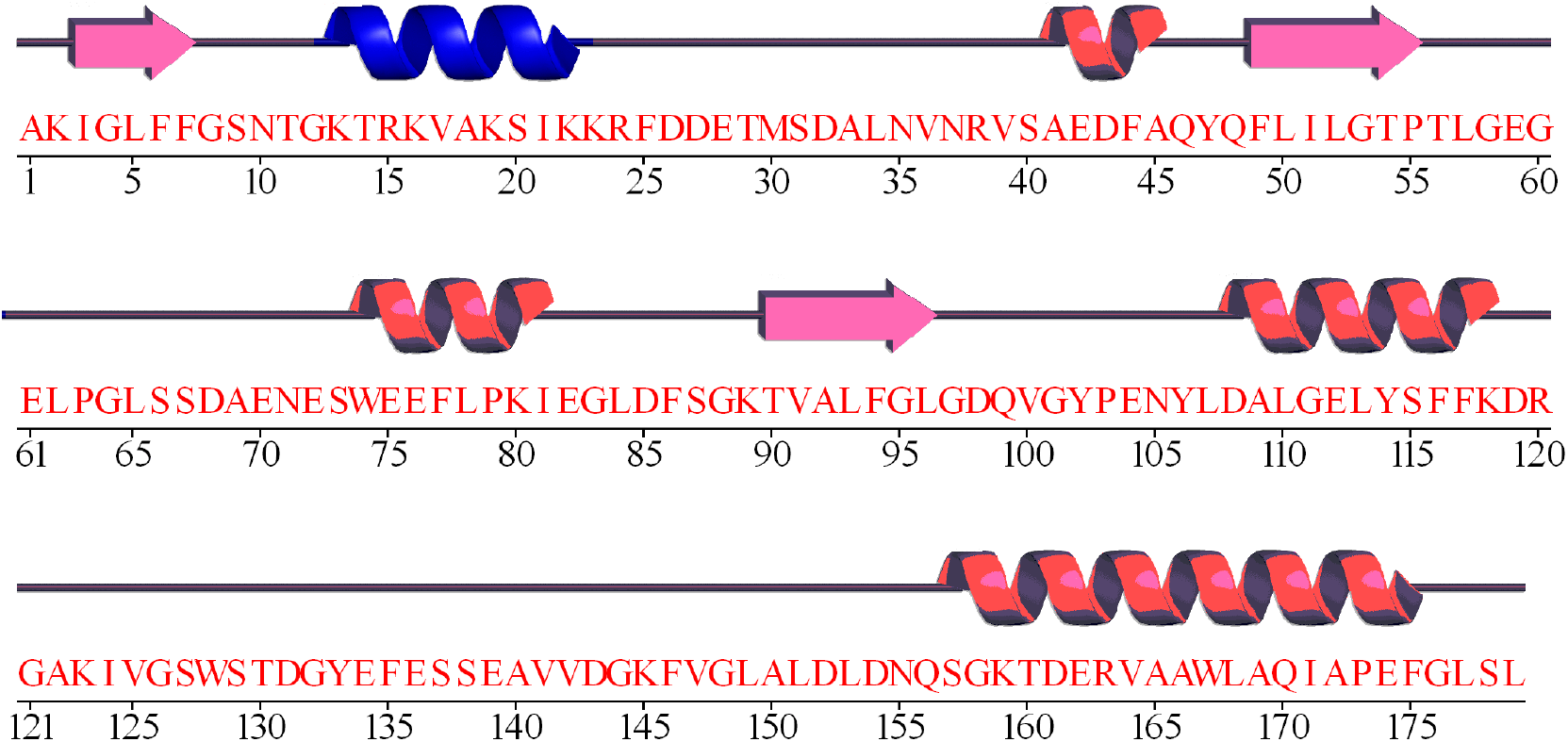

## D. COARSE-GRAINED POTENTIAL

We employed the following native-centric potential:

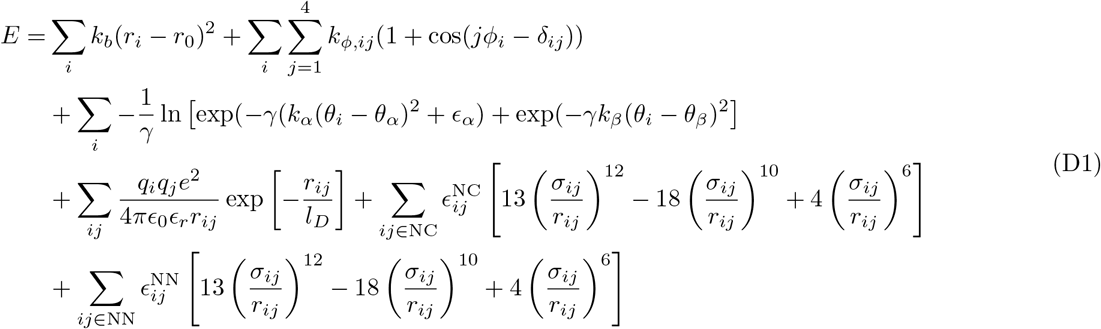

These terms are, in order, the contributions from bonds, dihedral angles (74), bond angles (75), electrostatics, native and non-native interactions (76). The bond term is a simple harmonic potential, where *k_b_* is the force constant, which is assigned to 100kJ/(*Å* · mol), *r*_0_ is the equilibrium bond length and *r_i_* is the actual distance between the beads. The dihedral angle term is the Karanicolas-Brooks potential (74), which corresponds to a standard periodic torsion potential where the force constants *k_ϕ,ij_*, and the phases *ϕ_ij_* are determined only by the second and third residue in every group of four residues defining a dihedral. The bond angle term is the Best-Hummer-Cheng potential (75), where *θ_i_* is the bond angle, and the parameters *γ, α, k_α_, k_β_* and *ϵ_α_* are constants (see (75) for more details).

The electrostatics term is a simple screened Coulombic potential with the charges defined by the net charge of the residues at pH=7 (*i.e*. lysine and arginine have a positive +1 charge, aspartate and glutamate have a negative −1 charge and the other residues are neutral). Finally, in the native and non-native potential, the value of 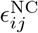, which sets the depth of the energy minimum for a native contact, is calculated as 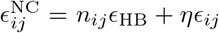. Here, *ϵ*_HB_, and *ϵ_ij_* represent energy contributions arising from hydrogen bonding and van der Waals contacts between residues *i* and *j* identified from the all-atom structure of the protein, respectively. *n_ij_* is the number of hydrogen bonds formed between residues *i* and *j* and *ϵ*_HB_ = 0.75 kcal/mol. The value of *ϵ_ij_* is set on the basis of the Betancourt-Thirumalai pairwise potential (77).

## E. ADDITIONAL FIGURES

**Figure S1.**
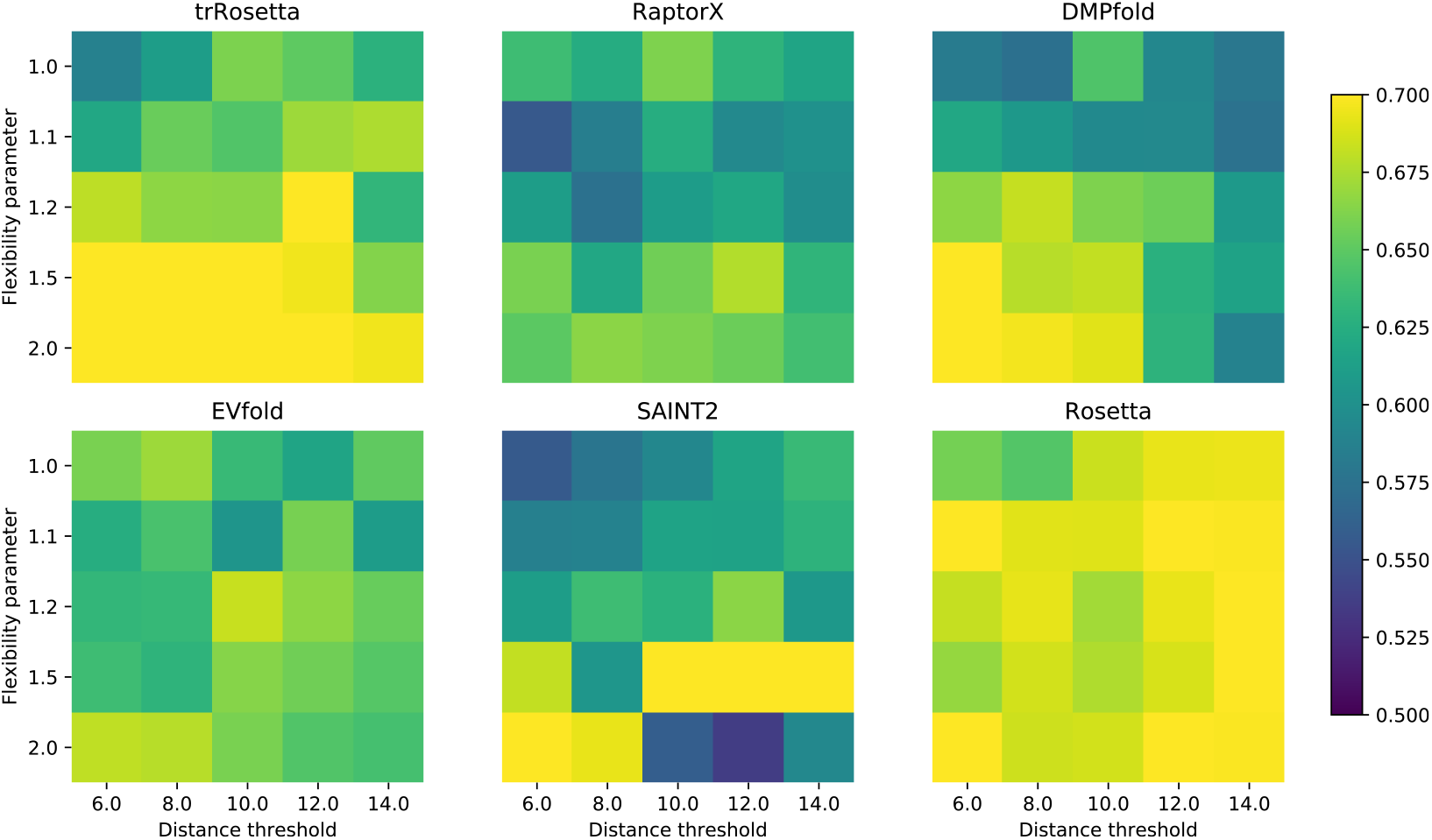
Parameter stability in trajectory analysis. We compare the area under the receiver-operating curve (AUROC) for the mechanism classification problem (determining if a protein folds via a two-state or multistate mechanism) for several choices of the distance threshold and flexibility hyperparameters. These results are produced using ten decoys per program for each of the 170 proteins presented in Appendix A. These numbers suggest that different parameter choices perform better for different programs, but there is little difference overall.

**Figure S2.**
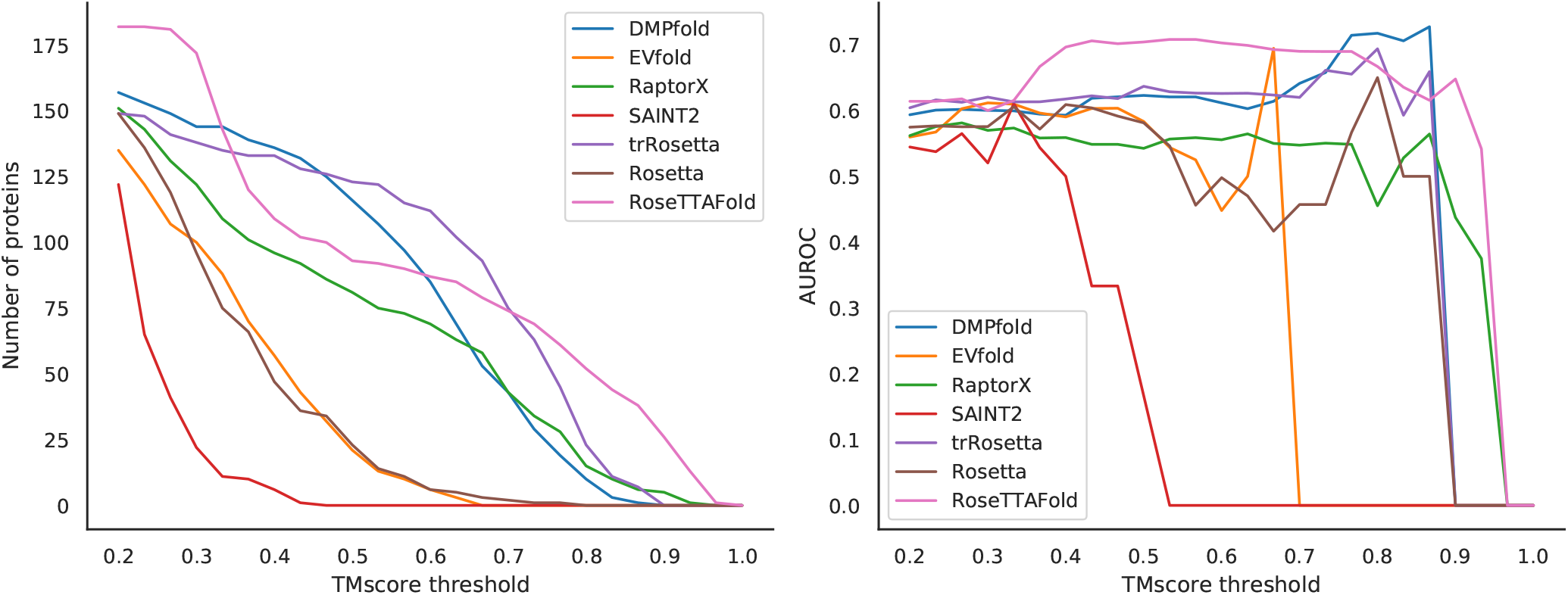
Predictive performance of the seven protein structure prediction programs. We compute the average TMscore (78) of the first ten decoys for each code, and use it as a proxy for the predictive performance of the algorithms. Left: cumulative number of proteins (*y*-axis) that were predicted with an average TMscore greater than the threshold (*x*-axis). The area under this curve can be interpreted as the global efficacy of the predictor. Right: area under the receiver-operator curve (*y*-axis) for all proteins above a given threshold (*x*-axis). For most proteins, the ability of simulated trajectories to distinguish formal folding kinetics is approximately independent from the predictive performance of the algorithm.

**Figure S3.**
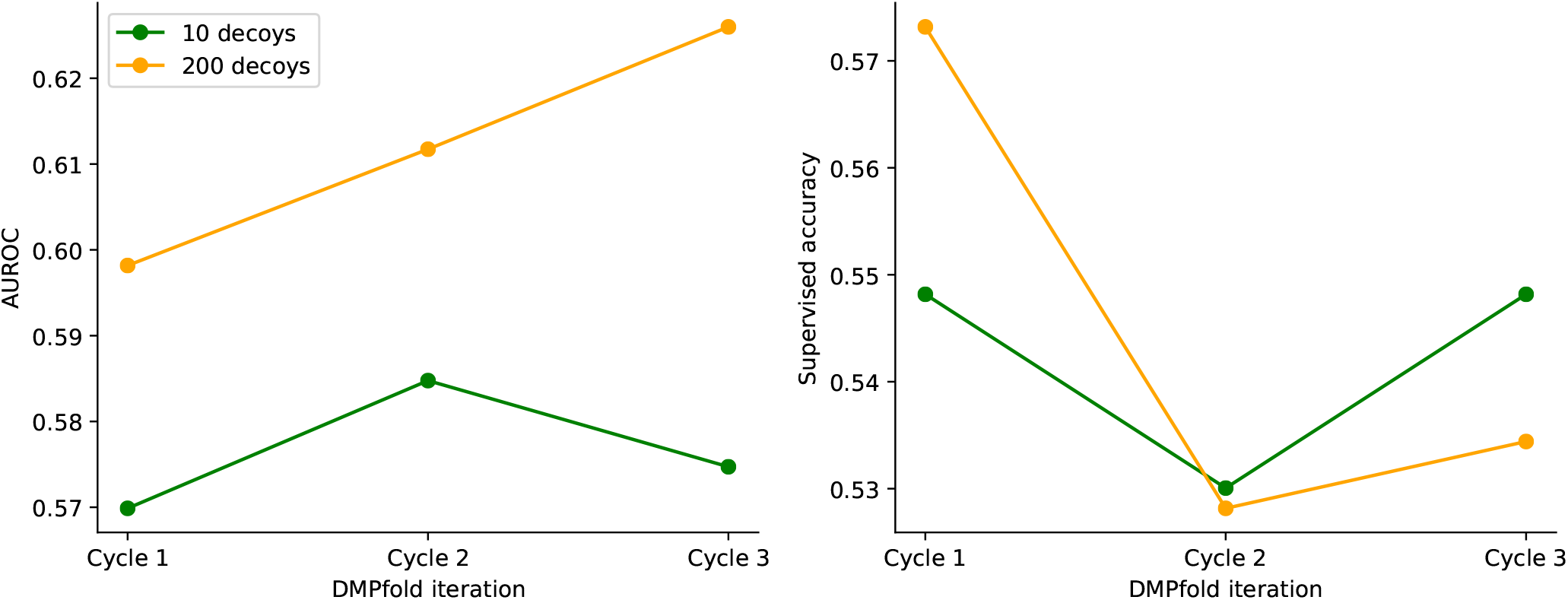
Change in predictive power across the different DMPfold cycles. While the reliability of the score, measured as the area under the receiver-operating curve (AUROC), seems to increase with successive cycles, the accuracy of the prediction does not improve.

**Figure S4.**
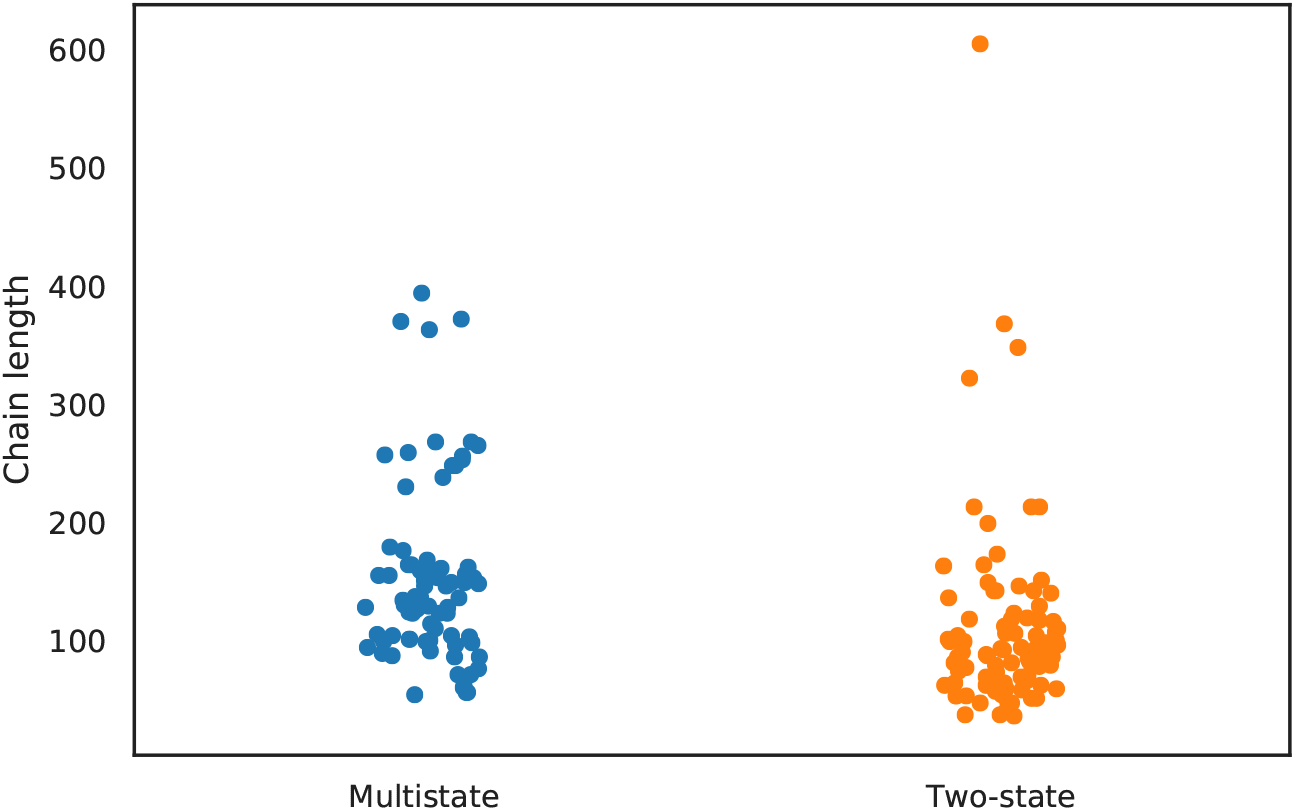
Distribution of lengths at each of the formal kinetics classes. There is not a trivial threshold that separates the classes.

**Figure S5.**
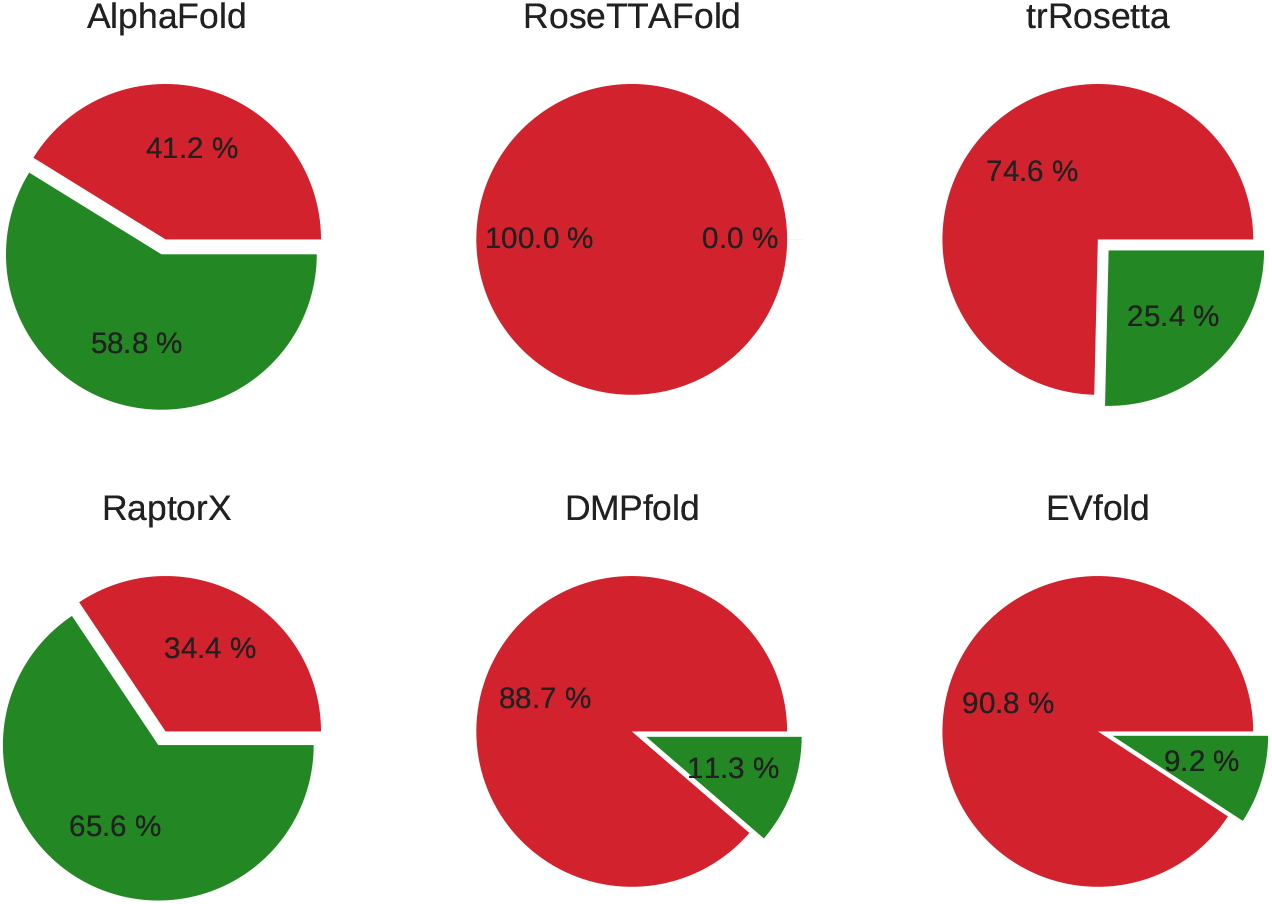
Mean proportion of the trajectories exhibiting significant clashes. A snapshot is considered to exhibit significant clashes if its clashscore (49) is higher than the 99th percentile for all PDB structures with resolution ≤ 2.5*Å* (30 in the snapshot downloaded on the 2nd of July of 2021).

**Figure S6.**
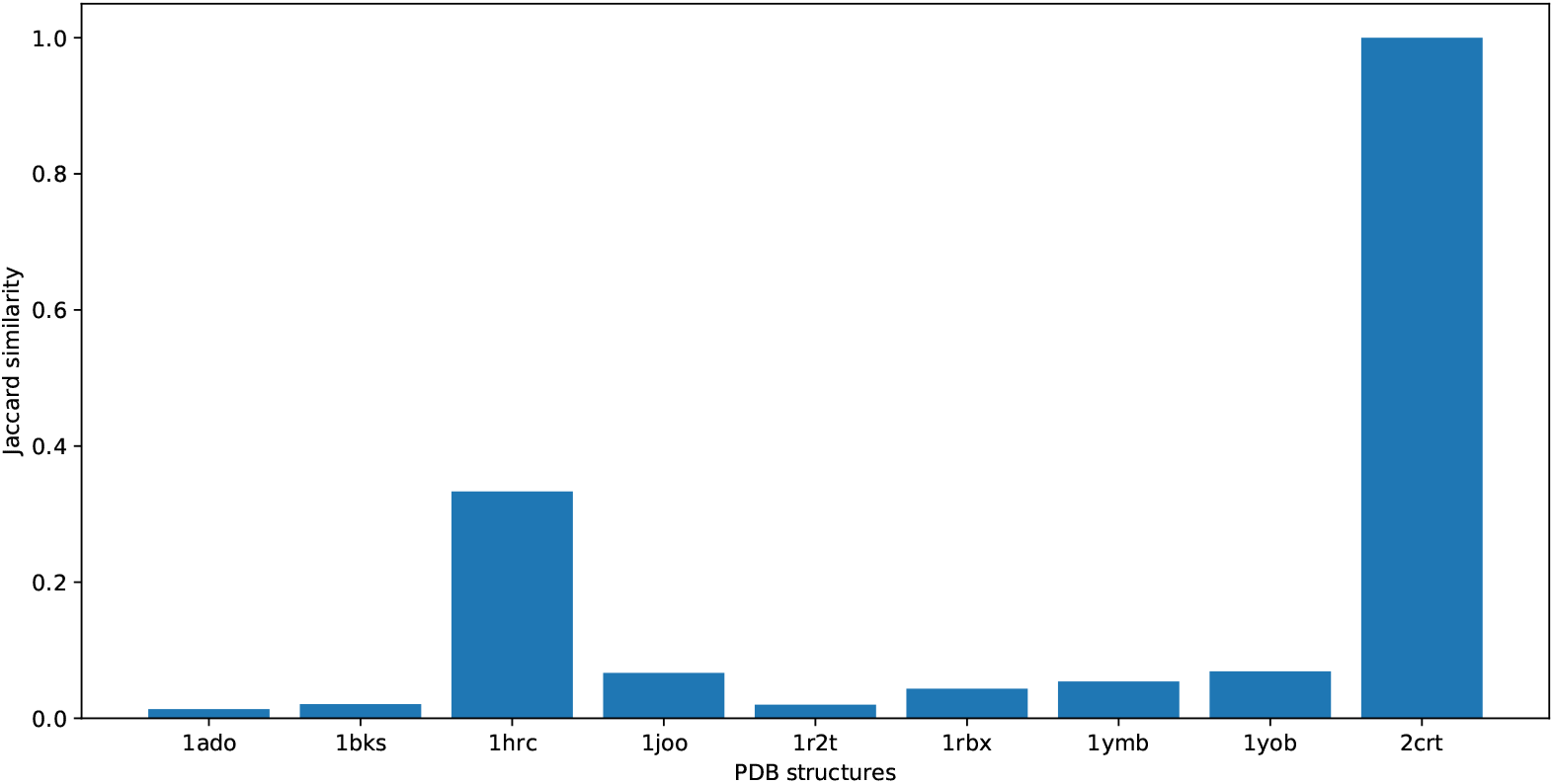
Jaccard similarity between the intermediates predicted by AlphaFold 2 and the ground truth. The assignments have been expressed as a binary string (where 1 means that the native contacts between secondary structure elements are formed in the intermediate, while 0 means they are not). AlphaFold 2 achieves a high score only on cardiotoxin analogue III (PDB: 2CRT), a small protein.

